# Quasi-periodic patterns contribute to functional connectivity in the brain

**DOI:** 10.1101/323162

**Authors:** Anzar Abbas, Michaël Belloy, Amrit Kashyap, Jacob Billings, Maysam Nezafati, Eric Schumacher, Shella Keilholz

## Abstract

Functional connectivity is widely used to study the coordination of activity between brain regions over time. Functional connectivity in the default mode and task positive networks is particularly important for normal brain function. However, the processes that give rise to functional connectivity in the brain are not fully understood. It has been postulated that low-frequency neural activity plays a key role in establishing the functional architecture of the brain. Quasi-periodic patterns (QPPs) are a reliably observable form of low-frequency neural activity that involve the default mode and task positive networks. Here, QPPs from resting-state and working memory task-performing individuals were acquired. The spatiotemporal pattern, strength, and frequency of the QPPs between the two groups were compared and the contribution of QPPs to functional connectivity in the brain was measured. In task-performing individuals, the spatiotemporal pattern of the QPP changes, particularly in task-relevant regions, and the QPP tends to occur with greater strength and frequency. Differences in the QPPs between the two groups could partially account for the variance in functional connectivity between resting-state and task-performing individuals. The QPPs contribute strongly to connectivity in the default mode and task positive networks and to the strength of anti-correlation seen between the two networks. Many of the connections affected by QPPs are also disrupted during several neurological disorders. These findings contribute to understanding the dynamic neural processes that give rise to functional connectivity in the brain and how they may be disrupted during disease.

**Highlights:**

- Quasi-periodic patterns (QPPs) of low-frequency activity contribute to functional connectivity
- The spatiotemporal pattern of QPPs differs between resting-state and task-performing individuals
- QPPs account for significant functional connectivity in the DMN and TPN during rest and task performance
- Changes in functional connectivity in these networks may reflect differences in QPPs

## 1 Introduction

Functional connectivity is a defining feature of resting-state functional magnetic resonance imaging (rs-fMRI). Correlation of the blood oxygenation level dependent (BOLD) signal fluctuations across brain regions is assumed to indicate coordinated activity between those regions (Biswal et al. 1995). Based on this assumption, maps of functional networks have been created from rs-fMRI data using multiple techniques (Park & Friston 2013; Power et al. 2011; Smith et al. 2013). The resulting functional networks agree closely with prevailing understandings of the functional organization of the brain (Asemi et al. 2015; Heuvel & Hulshoff Pol 2010; Vincent et al. 2008; Zhang et al. 2008). Consequently, functional connectivity has proven to be a useful tool in studying the brain, particularly when brain organization is disrupted during neurological disorders (Gillebert & Mantini 2013; for reviews, see Mohan et al. 2016; Pievani et al. 2014).

Despite the wide use of rs-fMRI and its clinical potential, the mechanisms that give rise to functional connectivity are not fully understood. In other words, we do not know what drives the coordination of neural activity in large-scale networks. It has been postulated that the functional architecture of the brain derives from low-frequency fluctuations of neural activity (Buzsáki 2006; Canolty & Knight 2010; He et al. 2008; Nir et al. 2008). In particular, infra-slow activity (< 1 Hz) has a similar frequency to BOLD fluctuations and appears highly relevant to functional connectivity between distant brain regions (Grooms et al. 2017; Hiltunen et al. 2014; Palva & Palva 2012; Pan et al. 2013). Phase-amplitude coupling between different frequencies of brain activity demonstrates the strong relationship between the infra-slow brain activity observable through BOLD and activity in higher frequency bands (Monto et al. 2008; Raichle et al. 2011; Thompson et al. 2014a). This suggests that neural dynamics in lower frequencies could provide a framework for the organization of functional systems (Foster et al. 2016).

Studies on functional connectivity typically focus on the average correlation between areas over the course of the scan. Several studies have shown that this approach ignores complex spatiotemporal patterns of activity such as global signal changes to propagating waves or time-lagged patterns of activation (Cole et al. 2016; Matsui et al. 2016; Mitra et al. 2016). Techniques for analysis of the dynamics of the BOLD signal allow for a more insightful understanding of real-time brain activity in rs-fMRI (Chang & Glover, 2010; for review, see Hutchison et al. 2013).

A quasi-periodic pattern (QPP) of large-scale network activity dominates BOLD fluctuations (Belloy et al. 2018; Majeed et al. 2011; Thompson et al. 2014b; Yousefi et al. 2018). It involves propagation of activity across several cortical and subcortical regions. Brain regions initially involved in the QPP are those within the default mode network (DMN). Activity then follows in regions pertaining to executive control, or the task positive network (TPN), alongside deactivation in the DMN. QPPs appear to reflect spatial patterns of infra-slow electrical activity (Pan et al. 2013; Thompson et al. 2014b; Grooms et al. 2017) and preliminary work shows that they may be linked to neuromodulation by deep brain nuclei (Abbas et al. 2018b). The wide spatial extent of the coordinated changes in the QPP is likely to contribute strongly to the BOLD correlation observed in the involved brain networks.

The DMN has been shown to exhibit altered connectivity in a variety of neurological and psychiatric disorders (for reviews, see Mohan et al. 2016; Raichle et al. 2015). It is possible that alterations in the QPP due to changes in neuromodulatory input could provide an economical explanation for the changes observed. Task performance drastically alters the functional architecture of the brain, shifting focus towards task-positive regions (Elton et al. 2015; Goparaju et al. 2014; Thompson et al. 2013). So far, a detailed analysis of QPPs has only been conducted in anesthetized animals and resting-state humans (Belloy et al. 2018; Majeed et al. 2009; Majeed et al. 2011; Thompson et al. 2014b; Yousefi et al. 2018). Task performance tends to increase anti-correlation between the DMN and TPN (Thompson et al. 2013), suggesting that QPP strength or frequency may be increased, altering measured functional connectivity as a result. The involvement of specific brain areas in any given task could also influence the spatiotemporal pattern of the observed QPPs.

In this study, we identified QPPs in humans during rest and while performing a working memory task. We looked for differences in the spatiotemporal pattern, strength, and frequency of the QPPs acquired from each group. We then minimized the QPPs’ contribution to the BOLD signal through a linear regression of the QPPs from the functional scans. By doing so, we could measure the impact of the QPP on functional connectivity by calculating functional connectivity before and after the regression. We looked at functional connectivity changes throughout the brain and specifically within and between the DMN and TPN. We hypothesized that removal of the QPPs from the functional scans through linear regression would lead to a reduction in functional connectivity strength within the DMN as well as a decrease in anti-correlation between the DMN and TPN. Our findings suggest that QPPs play an important role in maintaining the normal functional architecture of the two functional networks and that low-frequency activity in the form of QPPs contribute substantially to the organization of functional connectivity in the brain.

## 2 Methods

The Matlab script used for all analysis and figures included in this study is available on github.com/anzarabbas/qpps_rest_task.

### 2.1 Data acquisition and preprocessing

MRI data from 100 randomly-selected unrelated individuals (ages 22-36, 54 female) was downloaded from the Human Connectome Project (Van Essen et al. 2012). One anatomical scan was used for each individual (T1-weighted three-dimensional magnetization-prepared rapid gradient echo (T1w 3D MPRAGE) sequence; TR = 2400 ms, TE = 2.14 ms, TI = 1000 ms, FA = 8°, FOV = 224 mm × 224 mm, voxel size 0.7 mm isotropic) (Milchenko & Marcus 2012).

Two resting-state functional scans approximately 15 minutes in length were used (Gradient-echo Echo Planar Imaging; TR = 720 ms, TE = 33.1 ms, FA = 52°, FOV = 208 mm × 180 mm (RO × PE), matrix = 104 × 90 (RO × PE), slice thickness = 2.0 mm; 72 slices; 2.0 mm isotropic voxels, multi-band factor = 8, echo spacing 0.58 ms) with right-to-left (RL) phase encode direction in one scan and left-to-right (LR) phase encode direction in the other (Chen et al. 2015; Feinberg et al. 2010; Setsompop et al. 2011). Two working memory task functional scans were also used (RL and LR phase encode direction) with the same scan parameters as resting-state scans, except that the duration was approximately 5 minutes in length. Though any cognitively demanding task could have been chosen for this study, working memory was chosen for its high demand for controlled processing and its relatively long duration compared to other HCP task fMRI scans. To adjust for the difference in the lengths of resting-state and task-performing scans, the resting-state scans were truncated to the same length as the task-performing scans. The task, described in Barch et al. (2013), involved a version of the N-back task assessing working memory and cognitive control in block format. In each functional scan, there are 8 task blocks, each lasting 25 seconds, and 4 fixation blocks, each lasting 15 seconds. Half the task blocks use a 2-back working memory task whereas the other half use a 0-back working memory task. The blocks were divided into four categories; faces, places, tools, and body parts.

For all preprocessing steps, a combination of FSL 5.0 (Jenkinson et al. 2012) and MATLAB (Mathworks, Natick, MA) was utilized. First, anatomical data was registered to the 2 mm Montreal Neurological Institute (MNI) atlas using FLIRT (Jenkinson & Smith 2001; Jenkinson et al. 2002), brain-extracted using BET, and segmented into gray matter, white matter, and cerebrospinal fluid using FAST (Zhang et al. 2001). Functional data was motion corrected using MCFLIRT (Jenkinson et al. 2002), also registered to MNI space using FLIRT, spatially smoothed with a Gaussian kernel of 6 mm using FSLMATHS. Next, Matlab was used for a Fast Fourier Transform temporal filter with a bandpass between 0.01 Hz and 0.08 Hz; then the global, white matter, and CSF signals were regressed, and lastly, all voxel timecourses were z-scored.

### 2.2 Pattern acquisition

A spatiotemporal pattern-finding algorithm was used to search for repeating patterns of BOLD activation in the functional scans from resting-state and task-performing individuals separately. A detailed description of the algorithm used and the parameters inputted are outlined in Majeed et al. (2011). The pattern-finding algorithm selects a user-defined starting segment from within a functional scan and conducts a sliding correlation of the segment with the same functional scan. If the activity in the segment repeats at other instances in the functional scan, the resulting sliding correlation vector contains peaks indicating those occurrences. Additional segments are extracted at each of these instances and averaged together into an updated segment. Subsequent sliding correlations are then conducted between the continually updated segment and the functional scan. This process is repeated until the updated segment no longer shows variation and represents a reliably repeating pattern of activity within the functional scan. The result of the algorithm is a repeating spatiotemporal pattern from within the functional scan and a sliding correlation vector of the pattern with the functional scan itself.

There are two user-defined parameters that can influence the output of the algorithm: The length and location of the starting segment. As described in the Introduction, previous work in resting-state fMRI has shown that a reliably observable QPP lasts approximately 20 seconds. This has also been shown to be true for the resting-state and task-performing scans used in this study (Supplementary Figure 1; Supplementary Figure 2). Hence, a length of 20 seconds was used when selecting a starting segment for the pattern-finding algorithm. The location of the starting segment can affect the spatiotemporal pattern that is outputted by the algorithm. QPPs involve an initial increase in BOLD signal in regions within the DMN and a decrease in BOLD signal in regions within the TPN. The activity propagates along the cortex to a decrease in BOLD signal in the DMN and an increase in BOLD signal in the TPN. Simply put, the QPP consists of a propagation of BOLD activity between the DMN, or a DMN/TPN switch (Majeed et al. 2011; Yousefi et al. 2018). Though the pattern-finding algorithm has been shown to reliably output this pattern, the DMN/TPN switch can occur in varying phases depending on the location of the starting segment (Yousefi et al. 2018). To ensure the DMN/TPN switch occurs in the same phase in both groups, the algorithm is run multiple times for each group with starting segments selected at random locations in the functional scan.

For the resting-state and task-performing groups separately, 25 randomly-selected functional scans from unique individuals were concatenated. DMN and TPN maps for each group were acquired by selecting 10% of brain voxels most correlated and most anti-correlated with the posterior cingulate cortex respectively. For each group, the pattern-finding algorithm was applied to the concatenated functional scans 100 times with unique randomly-selected starting segments. All 100 patterns acquired for each group from the pattern-finding algorithm were analyzed for a DMN-to-TPN switch. The pattern most closely matching a DMN-to-TPN transition was selected and designated as a representative QPP for that group. By doing so, one representative QPP was chosen for the resting-state group and another for the task-performing group. The spatiotemporal pattern of the two QPPs was later compared. To demonstrate that the 25 individuals selected for QPP acquisition were not biasing the results, all pattern-acquisition steps were repeated with two subsequent groups of 25 randomly-selected scans from unique individuals and the results were compared across iterations (Supplementary Figure 3). For task-performing scans specifically, to demonstrate that the spatiotemporal pattern of the QPP did not differ between the first and second scans, all pattern-acquisition steps were repeated for the first and second scans separately and compared (Supplementary Figure 4).

Next, sliding correlation vectors were calculated between the two observed QPPs and functional scans from both resting-state and task-performing groups. Peaks, or local maxima, in the sliding correlation vectors signify occurrences of the QPP during the functional scan. This helps quantify the strength and frequency of QPP occurrence over time. The strength refers to correlation strength of the QPP with the functional scan, measured by the height of the peaks in the sliding correlation vectors. Frequency refers to how often these peaks occur. The sliding correlation vectors from all scans in each group were concatenated for the resting-state and task-performing QPPs separately. First, the mean height of peaks greater than 0.1 was calculated for the resting-state and task-performing QPPs in both the resting-state and task-performing scans. Second, the mean time interval between the peaks was also calculated for both QPPs in both groups. Third, the sliding correlation vectors were represented as histograms for comparison across groups without the need of the arbitrary 0.1 correlation threshold.

### 2.3 Blocks in the task-performing scans

As described earlier, the working memory task involved four 15-second fixation blocks, four 25-second 0-back working memory task blocks, and four 25-second 2-back working memory task blocks. To investigate the effects of the individual blocks on the QPPs, the sliding correlation vectors of the resting-state and task-performing QPPs were separated by block. The mean peak height for all peaks > 0.1 was calculated and the sliding correlation vectors were represented as histograms to be compared.

### 2.4 QPP regression

The resting-state and task-performing QPPs were regressed separately from the resting-state and task-performing scans to study their contributions to functional connectivity. For each functional scan, a unique regressor was calculated per brain voxel. This was done by convolving the QPP’s sliding correlation vector with the timecourse of each brain voxel during the QPP. The obtained regressor was z-scored to match the signal in the functional scan. Then, linear regression was carried out using standardized/beta coefficients and the regressors calculated for each brain voxel. This method produced a functional scan with attenuated presence of the QPP in the BOLD signal. The resting-state QPP was regressed from both resting-state and task-performing scans and the task-performing QPP was regressed from both resting-state and task-performing scans.

The efficacy of this regression method was demonstrated by conducting a subsequent sliding correlation of the QPPs with the QPP-regressed functional scans. Like the analysis of the concatenated sliding correlation vectors before QPP regression, the mean peak height and time interval between QPPs was calculated and the sliding correlation vectors were represented as histograms. The mean values and the histograms were compared to the ones created before QPP regression quantify the efficacy of the QPP regression in removing the presence of the QPPs in the functional scans.

### 2.5 Analysis of functional connectivity

A region of interest (ROI) atlas was used to summarize functional connectivity between all brain regions. Each functional scan was parceled into 273 ROIs from the Brainnetome Atlas (Fan et al. 2016). The mean signal over time, or timecourse, of each ROI was calculated. The ROI timecourses were used to acquire the strength of functional connectivity between brain regions through Pearson correlation. Functional connectivity strengths between all ROIs over the course of a functional scan were compiled into one functional connectivity matrix per scan. All functional connectivity matrices from each group underwent a Fischer’s z-transformation and were averaged into a mean functional connectivity matrix for that group. Functional connectivity matrices were also calculated for the functional scans after the QPPs had been regressed. Two-sample t-tests were performed for each ROI connection to check for significant differences in functional connectivity between groups. Multiple comparisons correction was performed by means of false discovery rate correction using the Benjamini and Yekutieli method (2001).

## 3 Results

### 3.1 Default mode and task positive networks

Maps for the default mode network and task positive network were acquired by locating areas strongly correlated or anti-correlated with the posterior cingulate cortex respectively (Figure 1a, 1b, *bottom-right panels*). The mean anti-correlation between the DMN and TPN in resting-state individuals was –0.78 with a standard deviation of 0.11. The mean anti-correlation between the DMN and TPN in task-performing individuals was –0.84 with a standard deviation of 0.10. The anti-correlation strength was significantly stronger in task-performing individuals with a *p* value of 4.78 × 10^-10^ calculated using a two-sample t-test.

**Figure 1:**
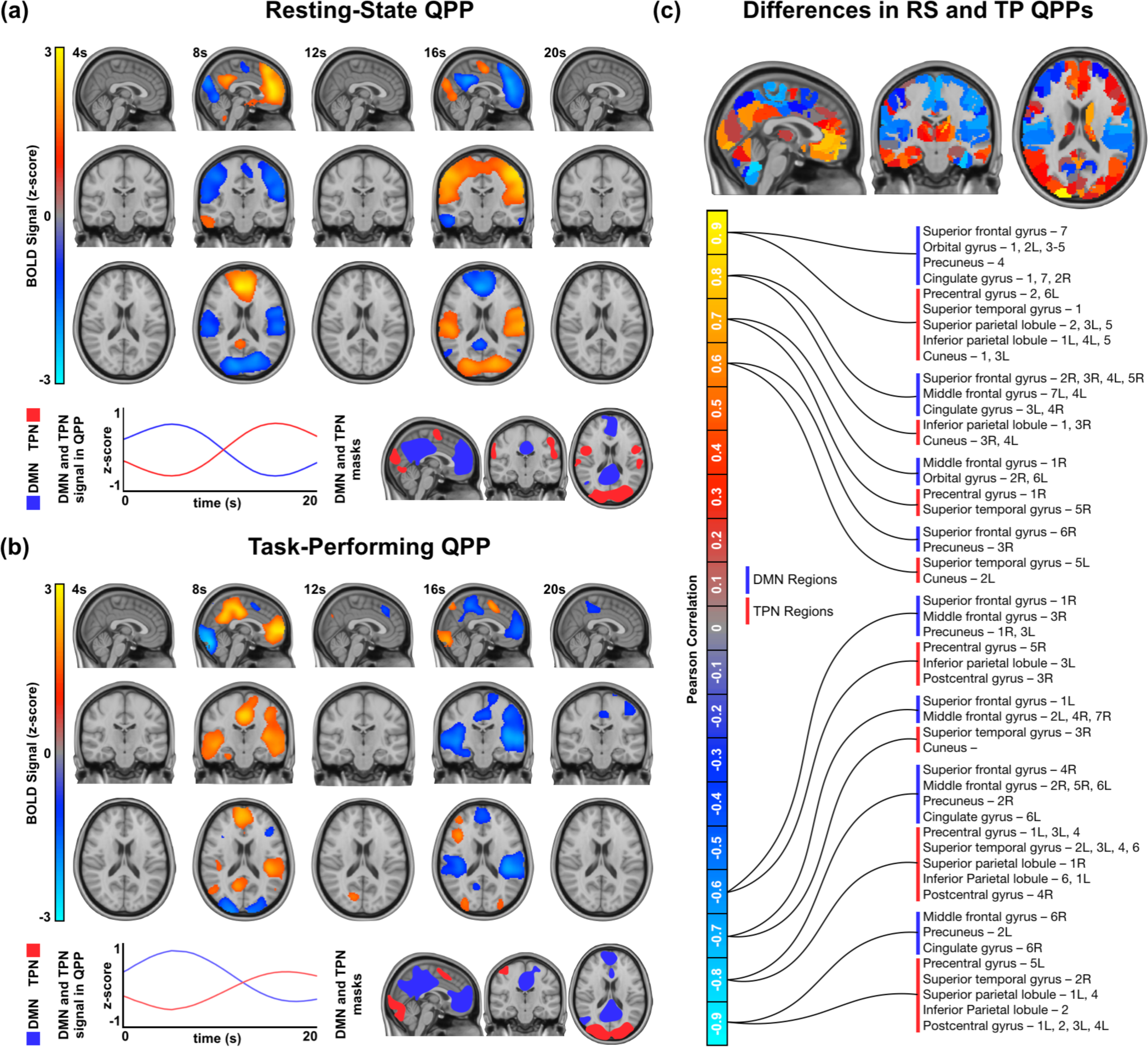
Quasi-periodic patterns in resting-state and task-performing groups. **(a)** *Top:* Spatiotemporal pattern seen in the resting-state QPP. Only BOLD signal changes 1.5x the standard deviation from the mean are shown. *Bottom-left*: DMN and TPN timecourse during the resting-state QPP. *Bottom-right*: Maps of DMN and TPN acquired from resting-state individuals. **(b)** *Top:* Spatiotemporal pattern seen in the task-performing QPP. Only BOLD signal changes 1.5x the standard deviation from the mean are shown. *Bottom-left*: DMN and TPN timecourses during the task-performing QPP. *Bottom-right*: Maps of DMN and TPN acquired from task-performing individuals. **(c)** Top: Spatiotemporal differences between the resting-state and task-performing QPPs. *Bottom:* Regions in the DMN and TPN that showed strong similarity between groups (> 0.6 Pearson correlation, shown in red/yellow) and strong dissimilarity between groups (< -0.6 Pearson correlation, shown in blue/turquoise). A full list of these regions can be found in Supplementary Table 1.

The default mode network map contained similar regions in both resting-state and task-performing individuals. For both the resting-state and task-performing groups, the DMN included parts of the superior and middle frontal gyri, orbital gyrus, paracentral lobule, middle and inferior temporal gyri, inferior parietal lobule, precuneus, cingulate gyrus, and cuneus. In the resting-state group, the DMN included parts of the cerebellum, which was not seen in the task-performing group. In the task-performing group, the DMN included parts of the precentral and postcentral gyri, superior temporal gyrus, superior parietal lobule, and striatum, which was not seen in the resting-state group.

The task positive network map also contained some variabilities between the resting-state and task-performing groups. For both groups, the TPN included parts of the superior and inferior frontal gyri, precentral and postcentral gyri, inferior temporal gyrus, fusiform gyrus, superior and inferior parietal lobules, insula, cuneus, occipital gyrus, and cerebellum. Unique to the resting-state group, the TPN included areas in the superior temporal gyrus. Unique to the task-performing group, the TPN included areas in the middle frontal gyrus.

### 3.2 Quasi-periodic patterns

#### 3.2.1 Comparison of spatiotemporal pattern

Application of the spatiotemporal pattern-finding algorithm resulted in the observation of a quasi-periodic pattern spanning 20 seconds in both resting-state and task-performing individuals (Figure 1a, 1b; Supplementary Figure 1; Supplementary Figure 2). QPPs acquired from application of the algorithm to 25 concatenated scans were representative of their respective groups (Supplementary Figure 3; Supplementary Figure 4). For both groups, the QPP involved an initial increase in BOLD signal in the DMN with decrease in BOLD signal in the TPN. This was followed by decrease in BOLD signal in the DMN and increase in BOLD signal in the TPN. Though DMN and TPN behavior was similar in both groups, there were differences in the specific brain regions involved.

A comparison of the spatiotemporal pattern between the two QPPs was conducted by comparing the mean activity of all ROIs during the course of the 20-second QPP. For each of the 273 ROIs, a Pearson correlation was conducted between its timecourse in the resting-state QPP and its timecourse in the task-performing QPP. Strong correlation signifies that the ROI behaved similarly in both groups, whereas a strong anti-correlation signifies the ROI behaved in the opposite manner. All ROIs within the DMN and TPN that were either strongly correlated (> 0.6 Pearson correlation) or strongly anti-correlated (< -0.6 Pearson correlation) are shown in Figure 1c. For the most part, DMN and TPN ROI timecourses were similar between the two groups, with differences dominated by task-relevant areas. Correlation strength of all 273 ROIs between the two groups’ QPPs can be found in Supplementary Table 1. Timecourses of example ROIs that were significantly different between the resting-state and task-performing QPPs are plotted in Supplementary Figure 5.

#### 3.2.2 Comparison of strength and frequency

Sliding correlation of both the resting-state and task-performing QPPs with all functional scans showed reliably recurring quasi-periodic peaks (Figure 2a, 2b), though at varying strengths and frequencies. By concatenating all sliding correlation vectors for each group, three comparisons of the temporal aspect of the QPPs were made:

**Figure 2:**
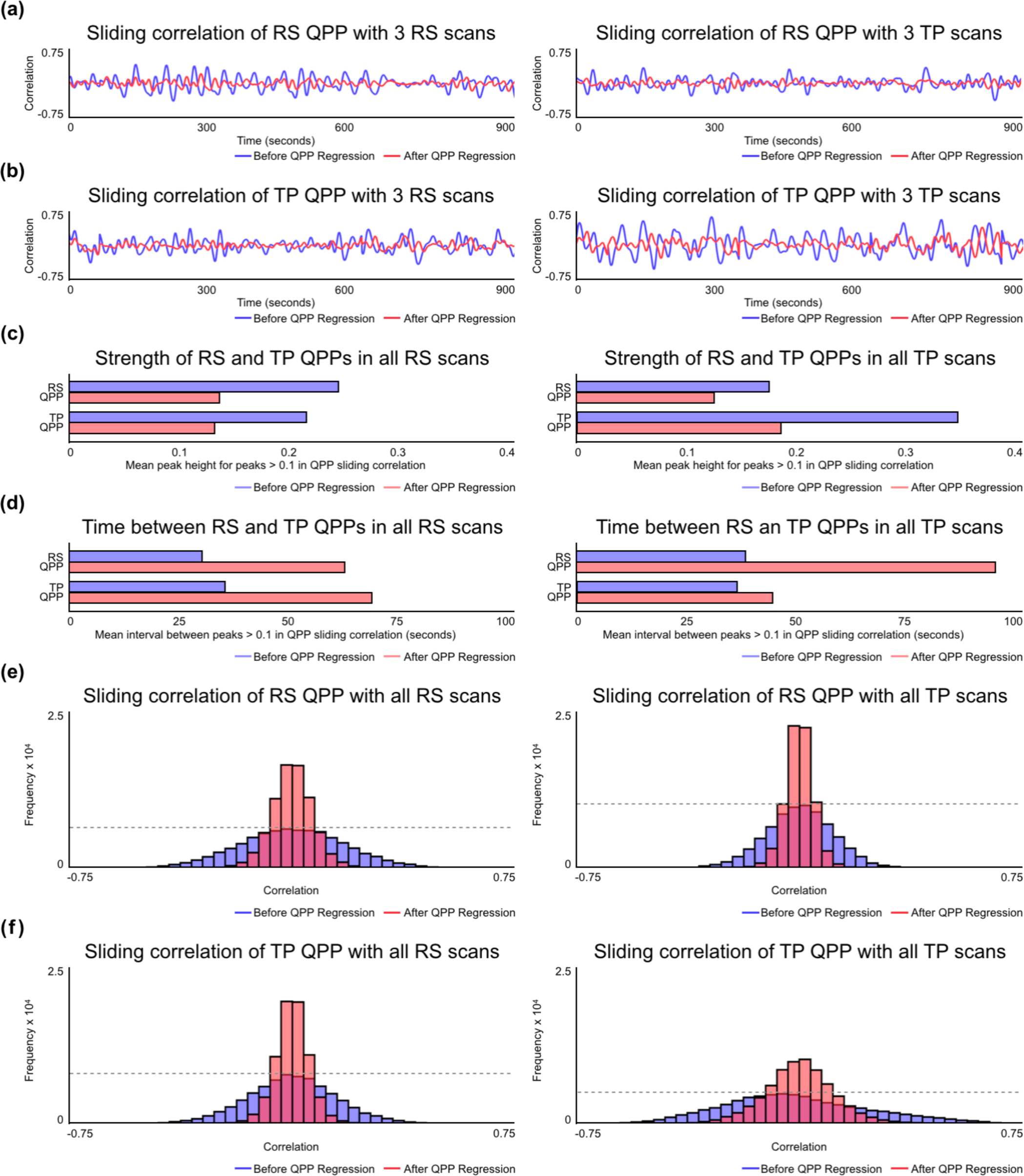
Strength and frequency of QPPs in resting-state and task-performing groups. (a) Example sliding correlation of the resting-state QPP with three concatenated scans from unique individuals during rest (left) and the same scans during task (right) before QPP regression (blue) and after QPP regression (red). (b) Example sliding correlation of the task-performing QPP with three concatenated scans from unique individuals during rest (left) and the same scans during task (right) before QPP regression (blue) and after QPP regression (red). (c) Mean correlation strength of peaks > 0.1 in the cumulative sliding correlation of the resting-state and task-performing QPPs with all resting-state scans (left) and all task-performing scans (right) before QPP regression (blue) and after QPP regression (red). (d) Mean time interval between peaks with correlation strength > 0.1 in the cumulative sliding correlation of the resting-state and task-performing QPPs with all resting-state scans (left) and all task-performing scans (right) before QPP regression (blue) and after QPP regression (red). (e) Histogram of the cumulative sliding correlation of the resting-state QPP with all resting-state scans (left) and all task-performing scans (right) before QPP regression (blue) and after QPP regression (red). (f) Histogram of the cumulative sliding correlation of the task-performing QPP with all resting-state scans (left) and all task-performing scans (right) before QPP regression (blue) and after QPP regression (red).

First, the mean correlation strengths at all peaks > 0.1 in the sliding correlation vectors were calculated and compared for the resting-state and task-performing QPPs in both groups (Fig 1c). The mean correlation strength of the resting-state QPP in resting-state scans was significantly higher than the task-performing QPP (*p* = 1.14 × 10^-08^). The mean correlation strength of the task-performing QPP in task-performing scans was significantly higher than the resting-state QPP (*p* = 1.7 x10^-110^). Also, the mean correlation strength of the task-performing QPP in task-performing scans was significantly higher than the resting-state QPP in resting-state scans (*p* = 5 × 10^-46^).

Second, the mean time intervals between all peaks > 0.1 in the sliding correlation vectors was calculated and compared for the resting-state and task-performing QPPs in both the resting-state and task-performing scans (Fig 1d). The mean time interval between resting-state QPP occurrences in the resting-state scans was significantly shorter than the task-performing QPP (*p* = 2.9 × 10^-09^). The mean time interval between task-performing QPP occurrences in the task-performing scans was shorter than the resting-state QPP, though with a relatively smaller significance (*p* = 0.0237). The mean time interval between task-performing QPP occurrences in the task-performing scans was significantly *longer* than the mean time interval between resting-state QPP occurrences in the resting-state scans (*p* = 9.8 × 10^-17^).

Third, the sliding correlation vectors of the resting-state and task-performing QPPs with all scans were represented as histograms for comparison between groups without the use of an arbitrary 0.1 correlation threshold (Fig 2e, 2f). A wide, short histogram indicates higher frequency of correlation values in the sliding correlation vector that are far from zero. This suggests a stronger presence of the QPP in the functional scan. A narrow, tall histogram indicates higher frequency of correlation values in the sliding correlation vector that are closer to zero. This suggests a weaker presence of the QPP in the functional scan. The histograms were compared using a Kolmogorov-Smirnov (KS) test and only significant differences with an alpha value of 1 × 10^-6^ are discussed in this paper. The resting-state QPP showed a stronger presence in the resting-state group compared to the task-performing group. Similarly, the task-performing QPP showed a stronger presence in the task-performing group compared to the resting-state group. Finally, the task-performing QPP showed a stronger presence in the task-performing group than the resting-state QPP showed in the resting-state group.

Additionally, the sliding correlation vectors of the resting-state and task-performing QPPs were compared for the fixation, 0-back, and 2-back task blocks in the working memory task scans. Comparison showed that the blocks did not have a significant effect on the strength and/or frequency of the QPPs (Supplementary Figure 6).

### 3.3 QPP regression

Linear regression was effective in attenuating the presence of QPPs in the functional scans. The sliding correlation vectors of the QPPs with QPP-regressed functional scans showed a diminished presence of the QPPs in the functional scans (Figure 2). For peaks in the sliding correlation vectors > 0.1, the mean correlation strength of the resting-state and task-performing QPPs with QPP-regressed resting-state and task-performing scans was significantly reduced (*p* = 1.3 × 10^-88^, *p* = 1.2 × 10^-58^, p = 1.2 × 10^-48^, p = 2.6 × 10^-92^ respectively). The mean time interval between occurrences of the resting-state and task-performing QPPs with QPP-regressed resting-state and task-performing scans significantly increased (*p* = 1 × 10^-33^, *p* = 3 × 10^-21^, *p* = 5.2 × 10^-26^, *p* = 1.4 × 10^-7^ respectively). The histograms representing the sliding correlation of the resting-state and task-performing QPPs with QPP-regressed resting-state and task-performing scans also showed a significantly weaker presence of the QPPs.

### 3.4 Overall functional connectivity differences

The functional connectivity matrices display the strength of functional connectivity in all 37,128 connections between the 273 ROIs in one image representing the static functional architecture of the brain. Data points closer to the central diagonal show functional connectivity strength in local connections while data points further away from the central diagonal show functional connectivity strength in long-range connections between brain regions.

An average functional connectivity matrix was calculated for resting-state and working memory task-performing individuals (Figure 3a). Significant functional connectivity differences between resting-state and task-performing individuals were widespread (Figure 3b, *bottom-left*), with 17,156 connections seeing a difference in functional connectivity strength. Native QPPs are those acquired from the same group; for example, the resting-state QPP is native to resting-state functional scans. Once native QPPs were regressed from all functional scans in each group, the number of functional connectivity differences between resting-state and task-performing individuals decreased by 40% to 10,259 (Figure 3b, *top-right*; Table 1).

**Figure 3:**
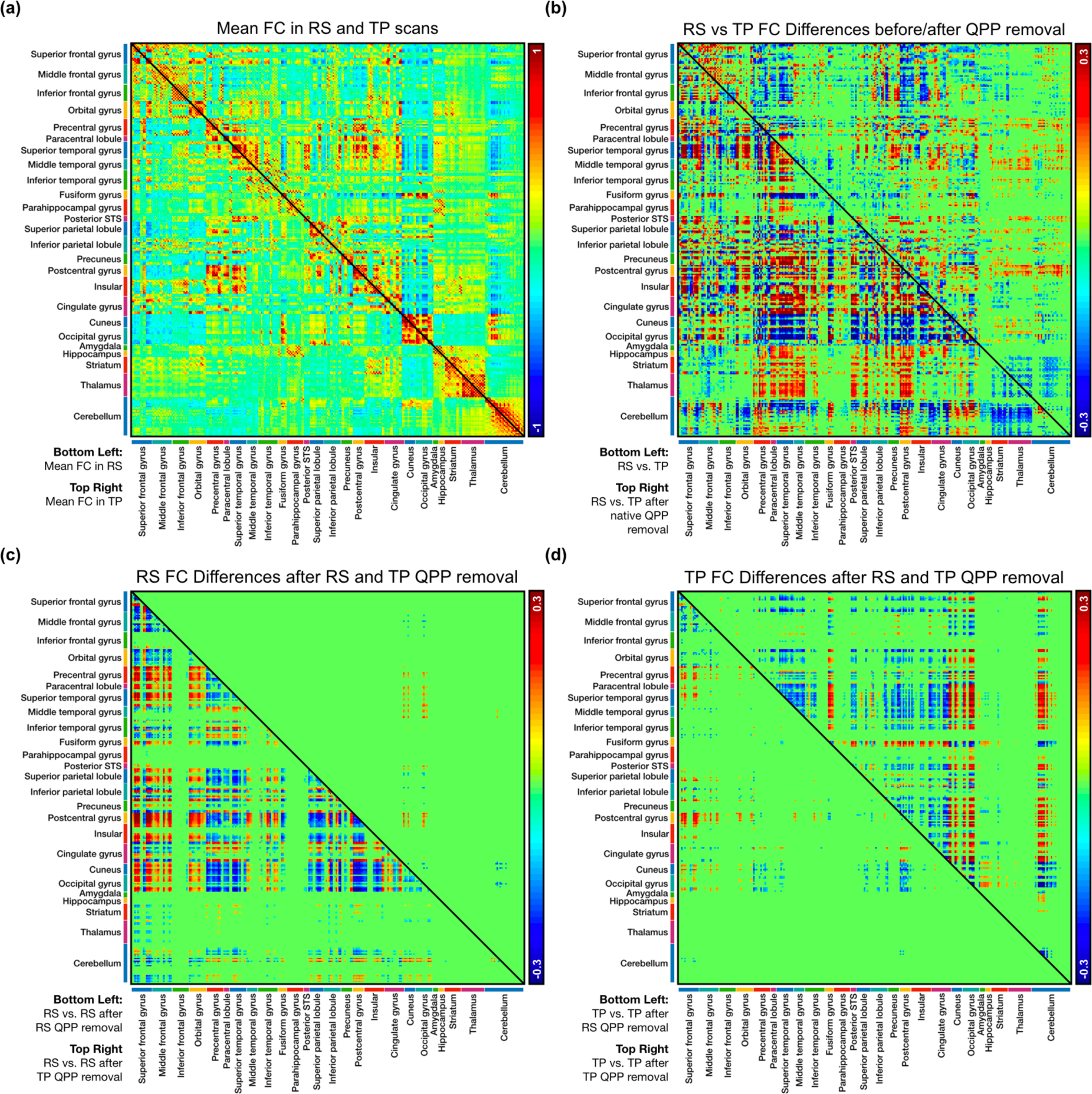
Functional connectivity (FC) in 273 regions of interest. **(a)** *Bottom-left*: Mean FC in the resting-state group. *Top-right*: Mean FC in the task-performing group. **(b)** *Bottom-left*: Significant differences in FC between the resting-state and task-performing group (*n* = 17,156). *Top-right*: Significant differences in FC between the resting-state and task-performing group after regression of their native QPPs (*n* = 10,259). **(c)** *Bottom-left*: Significant differences in FC in the resting-state group after regression of the resting-state QPP (*n* = 8,662). *Top-right*: Significant differences in FC in the resting-state group after regression of the task-performing QPP (*n* = 188). **(d)** *Bottom-left*: Significant differences in FC in the task-performing group after regression of the resting-state QPP (*n* = 1,062). *Top-right*: Significant differences in FC in the task-performing group after regression of the task-performing QPP (*n* = 5,756).

**Table 1:** Description of functional connectivity differences between the resting-state and task-performing groups before and after regression of their native QPPs. When comparing significant functional connectivity differences between the original functional scans and after the QPPs had been regressed, the first column shows the percent distribution of the different directions the functional connectivity changes occurred in, the second column shows the mean magnitude shift in strength of Pearson correlation for each of the directions, and the third column shows the total number of ROI connections with a significant change in functional connectivity between groups. The total number of significant changes in functional connectivity decreased by 40% after regression of native QPPs.

|  | FC changes |  | Original functional scans |  |  | After regression of native QPPs |  |  |
| --- | --- | --- | --- | --- | --- | --- | --- | --- |
| Resting-state vs. Task-performing | Increases | – → + | 44% | ↑ 0.24 | 9,133 | 34% | ↑ 0.17 | 5,138 |
|  |  | + → + | 27% | ↑ 0.19 |  | 28% | ↑ 0.17 |  |
|  |  | – → – | 29% | ↑ 0.17 |  | 38% | ↑ 0.14 |  |
|  | Decreases | + → – | 33% | ↓ 0.23 | 8,023 | 31% | ↓ 0.17 | 5,121 |
|  |  | – → – | 42% | ↓ 0.20 |  | 41% | ↓ 0.15 |  |
|  |  | + → + | 25% | ↓ 0.18 |  | 28% | ↓ 0.16 |  |

Regression of the resting-state QPP from resting-state functional scans led to 8,662 significant changes in functional connectivity (Figure 3c, *bottom-left*). When the task-performing QPP was regressed from the resting-state scans, only 188 connections were significantly altered (Figure 3c, *top-right*; Table 2). Regression of the task-performing QPP from task-performing functional scans led to 5,756 significant changes in functional connectivity (Figure 3d, *top right*). When the resting-state QPP was regressed from the task-performing scans, the number of significant changes decreased to 1,062 (Figure 3d, *bottom-left*; Table 2).

**Table 2:** Description of functional connectivity changes in the resting-state and task-performing groups after regression of the resting-state QPP and after regression of the task-performing QPP. For the resting-state and task-performing groups separately, when comparing significant changes in functional connectivity after regression of the resting-state QPP and after regression of the task-performing QPP, the first column shows the percent distribution of the different directions the functional connectivity changes occurred in, the second column shows the mean magnitude shift in strength of Pearson correlation for each of the directions, and the third column shows the total number of ROI connections with a significant change in functional connectivity between groups. Regression of the task-performing QPP from the resting-state scans showed a 98% decrease in significant functional connectivity changes when compared to regression of the resting-state QPP form resting-state scans. Regression of the resting-state QPP from the task-performing scans showed an 82% decrease in significant functional connectivity changes when compared to regression of the task-performing QPP from task-performing scans.

|  | FC changes |  | Original functional scans |  |  | After regression of native QPPs |  |  |
| --- | --- | --- | --- | --- | --- | --- | --- | --- |
| Resting-state | Increases | - → + | 22% | ↑ 0.18 | 4,271 | 60% | ↑ 0.15 | 139 |
|  |  | + → + | 7% | ↑ 0.14 |  | 27% | ↑ 0.15 |  |
|  |  | - → - | 71% | ↑ 0.16 |  | 13% | ↑ 0.13 |  |
|  | Decreases | + → - | 35% | ↓ 0.18 | 4,391 | 26% | ↓ 0.14 | 49 |
|  |  | - → - | 21% | ↓ 0.15 |  | 33% | ↓ 0.14 |  |
|  |  | + → + | 44% | ↓ 0.16 |  | 41% | ↓ 0.16 |  |
| Task-performing | Increases | - → + | 26% | ↑ 0.16 | 553 | 8% | ↑ 0.15 | 2,481 |
|  |  | + → + | 27% | ↑ 0.16 |  | 2% | ↑ 0.13 |  |
|  |  | - → - | 47% | ↑ 0.14 |  | 90% | ↑ 0.19 |  |
|  | Decreases | + → - | 18% | ↓ 0.15 | 508 | 29% | ↓ 0.16 | 3,275 |
|  |  | - → - | 61% | ↓ 0.15 |  | 6% | ↓ 0.13 |  |
|  |  | + → + | 21% | ↓ 0.15 |  | 65% | ↓ 0.17 |  |

### 3.5 Functional connectivity changes in the DMN and TPN

QPP regression particularly affected connections within the DMN and TPN as well as between the two networks. After regression of the resting-state QPP from resting-state scans, there was a strong decrease in local functional connectivity in the anterior regions of the DMN, namely the superior frontal, middle frontal, and orbital gyri. There was also a sharp decrease in connectivity between the anterior and posterior regions of the DMN. Additionally, the anti-correlation between the DMN and TPN diminished significantly. The TPN itself showed a decrease in functional connectivity, both locally and across regions (Figure 4a, bottom-left). Alternatively, regression of the task-performing QPP from resting-state scans did not result in as widespread changes in the DMN and TPN (Figure 4a, top-right).

**Figure 4:**
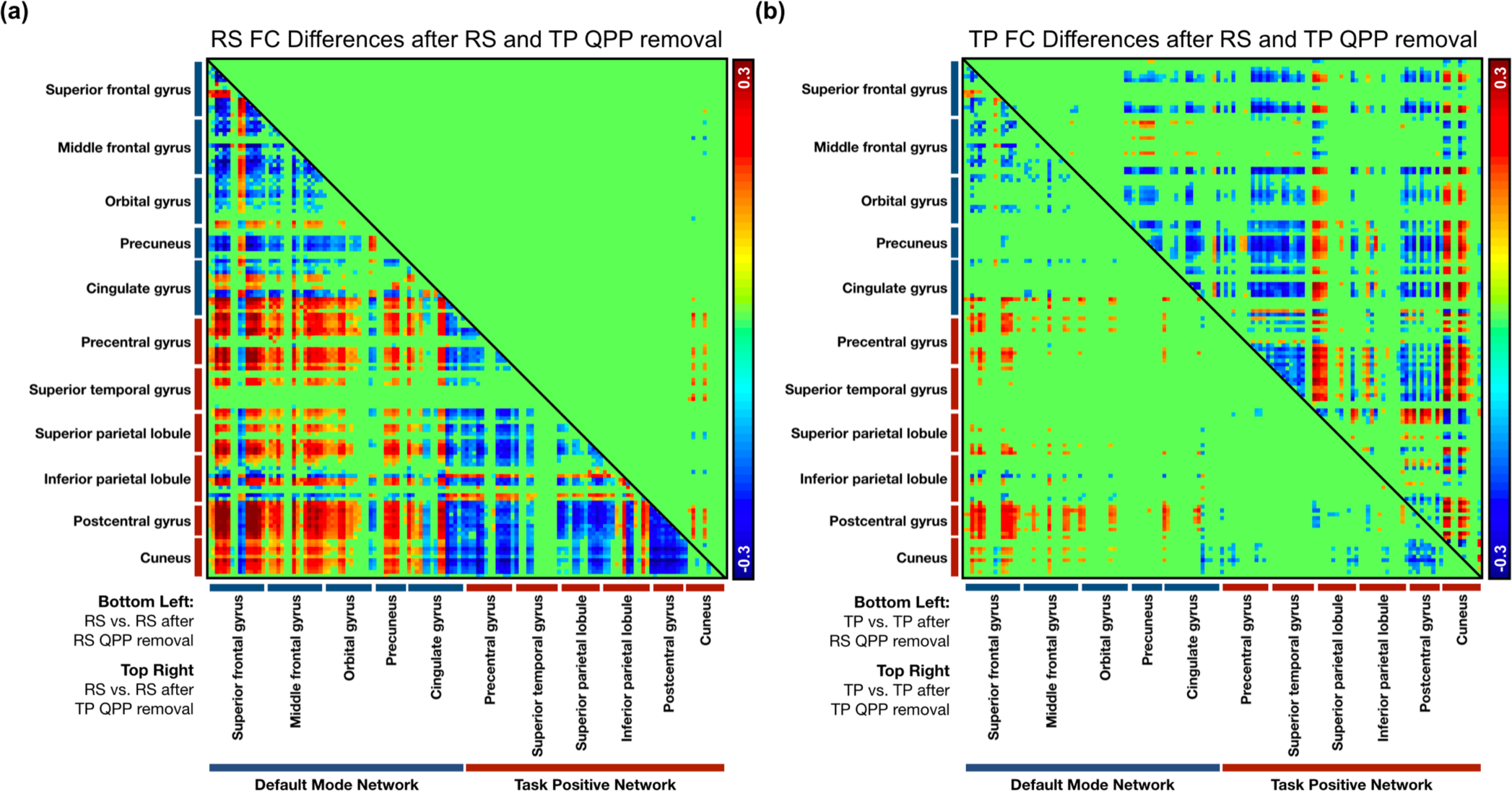
Significant functional connectivity changes in regions within the DMN and TPN after regression of QPPs. **(a)** *Bottom-left*: Significant differences in FC in the resting-state group after regression of the resting-state QPP. *Top-right*: Significant differences in FC in the resting-state group after regression of the task-performing QPP. **(b)** *Bottom-left*: Significant differences in FC in the task-performing group after regression of the resting-state QPP. *Top-right*: Significant differences in FC in the task-performing group after regression of the task-performing QPP.

Regression of the task-performing QPP from task-performing scans also affected areas in the DMN and TPN (Figure 4b, top-right). Similar to the resting-state group, there was a decrease in functional connectivity between anterior and posterior regions of the DMN. However, the local decreases in functional connectivity were seen in the posterior regions, namely the precuneus and cingulate gyrus. There were both decreases and increases in anti-correlation between the DMN and TPN, and a mixture of decreases and increases in functional connectivity between regions within the TPN. The functional connectivity changes when the resting-state QPP was regressed from the task-performing group showed similarity with resting-state individuals, though were significantly smaller in number.

## 4 Discussion

Our study reproduces previous reports of the presence of reliably recurring quasi-periodic patterns in the brain (Belloy et al. 2018; Majeed et al. 2011; Thompson et al. 2014; Yousefi et al. 2018). This is the first time QPPs have been examined in whole-brain data from task-performing individuals. Comparison of the QPPs acquired from resting-state and task-performing individuals show distinct spatiotemporal differences. These differences are specific to brain regions involved in the working memory task, suggesting that variability in the QPPs may be task-specific. Regression of the QPPs from the functional scans showed that QPPs have a strong effect on connectivity strength between brain regions. This effect is concentrated towards DMN and TPN functional connectivity, two networks central to the pattern.

### 4.1 DMN and TPN differences across groups

Details on the working memory task used in this study can be found in Barch et al. (2013), along with the brain regions activated and deactivated during the 0-back and 2-back tasks respectively. Overall, presence of a task led to strong activations in TPN regions and deactivations in DMN regions. Since the DMN and TPN maps for resting-state and task-performing scans were created using all time points irrespective of task blocks, the differences in brain regions that were involved in either task block are not considered when comparing DMN and TPN maps between the groups.

Regions mapped as DMN areas in this study largely agreed with previous findings (Fox et al. 2005). However, there were some differences in DMN maps created from the resting-state group and the task-performing group. The inclusion of certain cerebellar regions in the DMN was unique to the resting-state group. These same regions were activated during task (Barch et al. 2013), reducing their co-activation with other DMN regions in the task-performing group. Inclusion of parts of the precentral and postcentral gyri, superior temporal gyrus, superior parietal lobule, and striatum in the DMN was unique to the task-performing group. The precentral and postcentral gyri as well as the superior temporal gyrus were deactivated during task. This increase in anti-correlation with TPN areas is likely why these regions were included in the DMN map in the task-performing group. The superior parietal lobule and striatum did not follow this pattern as they were activated during task.

The TPN map also agreed with previous findings (Fox et al. 2005) and was similar across groups, with exceptions. A small area in the middle frontal gyrus was included in the TPN map for task-performing individuals. The middle frontal gyrus is one of the regions strongly activated during task, when the DMN is being deactivated. Its categorization as a TPN area unique to task-performing individuals is likely due to this anti-correlation with the DMN during the task. Similarly, inclusion of the superior temporal gyrus in the TPN was unique to the resting-state group. This region showed a strong deactivation during task. This lack of anti-correlation with DMN regions is likely why the superior temporal gyrus was not part of the TPN in the task-performing group.

### 4.2 Spatiotemporal pattern of QPPs

A spatiotemporal comparison of the DMN portion of the QPPs acquired from both groups demonstrates differences similar to those noted in the DMN maps. Both groups’ QPPs show a strong increase in BOLD signal in the DMN in the first half of the pattern followed by a decrease in BOLD signal in DMN regions in the second half. As a result, DMN regions are highly correlated between the two groups’ QPPs. However, there are exceptions. This includes regions in the superior and middle frontal gyri, the cingulate gyrus, precuneus, inferior parietal lobule, paracentral lobule, precentral gyrus, and superior parietal lobule. These regions follow the opposite trend in the task-performing QPP when compared to the resting-state QPP. This is consistent with the observation that some of these regions are only seen in the DMN map from the task-performing group, and not the resting-state group. Both groups’ QPPs also show a strong decrease in BOLD signal in the TPN in the first half of the pattern followed by an increase in BOLD signal in TPN regions in the second half. As was seen with the DMN, there are exceptions. This includes regions in the superior and middle frontal gyri, the superior and middle temporal gyri, the pre-and post-central gyri, superior and inferior parietal lobule, fusiform gyrus, insula, and cerebellum. These regions follow the opposite trend in the task-performing QPP when compared to the resting-state QPP. Interestingly, Barch et al. (2013) shows that many of the regions that behave differently across the two QPPs are also relevant to the task being performed.

The robust spatiotemporal differences in the QPPs during task performance compared to resting-state are intriguing and suggest that the QPP is not a fixed pattern of coordinated activity but rather a flexible framework that organizes the brain into large-scale networks to optimize task performance. Thus, variabilities in the QPP’s spatial pattern may be task-specific, and other tasks involving different brain regions may alter the spatial pattern of the QPP in a different way.

Since application of the pattern-finding algorithm requires a few minutes of continuous data at the very least, spatiotemporal comparison of the QPPs across the fixation, 0-back, and 2-back blocks during the task-performing scan is not feasible. Given that all time points in the scans were used to acquire QPPs, the differences highlighted between the resting-state and task-performing QPPs are meant to reflect changes as an overall product of task performance rather than changes specifically due to 0-back or 2-back tasks.

### 4.3 Strength and frequency of QPPs

The QPPs acquired from both the resting-state and task-performing groups showed quasi-periodic peaks in their sliding correlation vectors with all functional scans. Unsurprisingly, the resting-state and task-performing QPPs showed stronger presence in their native scans compared to the opposing scans. Additionally, the task-performing QPP showed greater correlation strength in the task-performing scans than the resting-state QPP did in resting-state scans, which may account for the significantly stronger anti-correlation between the DMN and TPN in the task-performing group observed here and in prior studies (Hampson et al. 2010; Kelly et al. 2008).

Though a spatiotemporal comparison of the QPPs between fixation, 0-back, and 2-back blocks was not possible, a comparison of the strength and frequency of the QPPs in each task block was conducted. The sliding correlation vectors of both resting-state and task-performing QPPs did not show any differences across the three blocks in the task-performing scans. Hence, the differences in the strength and frequency of QPPs highlighted between resting-state and task-performing individuals are meant to highlight changes as an overall result of task performance, rather than an effect of 0-back or 2-back tasks specifically.

A cognitively demanding task such as the one used in this study leads to an inherently higher state of vigilance compared to rest. Given the QPP’s involvement of both DMN and TPN activity, its possible origin in neuromodulatory input, and the relationship between infra-slow electrical activity and vigilance levels, it seems plausible that the greater strength of the QPP in the task-performing group might arise from the increased alertness needed for task performance. A preliminary analysis conducted in data from rhesus macaques showed that QPPs occur with greater strength and frequency in awake macaques compared to anesthetized macaques (Abbas et al. 2016), suggesting that vigilance may be playing a role. Another study showed that when performing a psychomotor vigilance task, greater anti-correlation between the DMN and TPN was tied to faster performance on the task (Thompson et al. 2013), which may be tied to the strength of the QPP.

The observation of the resting-state QPP in task-performing scans and the task-performing QPP in resting-state scans is likely due to the similarity in the spatial patterns of the QPPs. As described above, though there are distinct differences in the QPPs acquired from both groups, there are strong similarities as well. When a sliding correlation is being calculated, any similarities between the resting-state and task-performing QPPs may lead to higher correlation values than if the spatial pattern of the QPPs had been entirely different. If this is true, then the sliding correlation vectors of the resting-state and task-performing QPPs should align for the same scan. To investigate this, a cross-correlation of the sliding correlation vectors of the resting-state QPP and task-performing QPP in both groups was conducted. At the lag showing maximum correlation strength, the correlation between the two QPPs’ sliding correlation vectors was 0.55. This suggests that the sliding correlation vectors of the two QPPs could indeed be aligned, which would explain the ‘presence’ of the resting-state QPP in task-performing scans and vice versa.

### 4.4 Functional connectivity changes

With regression of the resting-state QPP from resting-state scans, there were strong local functional connectivity decreases in the anterior regions of the DMN and in the longer-range connections between anterior and posterior DMN nodes. A general decrease in connectivity within the TPN also occurred. As would be expected, there was an attenuation of anti-correlation between the DMN and TPN. Loss of functional connectivity in these regions is significant in that many neurological and psychiatric patients exhibit similar connectivity disruptions during resting-state scans. Previous studies have shown that local connectivity changes in the anterior regions of the DMN are associated with schizophrenia (Holt et al. 2011), while reduced functional connectivity between anterior and posterior regions of the DMN in associated with ADHD (Choi et al. 2013), Alzheimer’s Disease (Sheline et al. 2010a; Zhang et al. 2009), as well as aging (Andrews-Hanna et al. 2007). The relationship between DMN and TPN activity and the strength of their anti-correlation is important for normal brain function and task performance (Fox et al. 2005; Thompson et al. 2013). Decreased anti-correlation between the DMN and TPN is even seen in individuals with ADHD (Hoekzema et al. 2013; Posner et al. 2014) and is restored after treatment with atomoxetine and methylphenidate (Liddle et al. 2010; Lin et al. 2015). A preliminary analysis by our group has also shown that QPPs are disrupted in individuals with ADHD (Abbas et al. 2018a). Given that regression of the QPP leads to altered functional connectivity in those same connections, a disruption of the QPP may be one of the factors in the development of such disorders.

Regression of the task-performing QPP from resting-state scans did not result in many significant changes in functional connectivity. This suggests that the task-performing QPP may be particular to the task or task-performing states in general, and may occur only rarely while the individual is at rest.

In task-performing scans, functional connectivity differences were seen after regression of both the resting-state and task-performing QPPs. Once more, the differences seen pertained to DMN and TPN regions and their interconnectivity. Similar to regression of the resting-state QPP from resting-state scans, regression of the task-performing QPP from task-performing scans led to an overall decrease in functional connectivity strength between anterior and posterior regions of the DMN. Unlike the resting-state group, the major short-range functional connectivity decreases in the DMN were seen in its posterior node. Additionally, regression of the task-performing QPP in task-performing scans led to a decrease in functional connectivity between DMN and TPN areas. Since the DMN and TPN are already anti-correlated, this meant a stronger anti-correlation between the two networks. This suggests that though the resting-state QPP may be playing a constructive role in distinguishing DMN and TPN regions from each other, the task-performing QPP in task-performing scans is doing the opposite. Hyper-connectivity between anterior and posterior regions of the DMN is seen during Major Depressive Disorder particularly during task performance as a potential result of the inability to shut off DMN activity during tasks (Grimm et al. 2008; Sheline et al. 2009; Sheline et al. 2010b). There were two regions that showed the similar changes to functional connectivity as the resting-state group, namely the superior parietal lobule and the cuneus. However, even for these regions, the middle frontal gyrus showed an increase in anti-correlation rather than a decrease.

Regression of the resting-state QPP from task-performing scans did result in some significant differences in functional connectivity. These changes followed the same trend as when the resting-state QPP was regressed from resting-state scans, albeit at a much smaller scale. These findings suggest that the resting-state QPP may still be occurring at a weaker frequency in the task-performing state. If so, it would be serving a similar purpose in maintaining strong functional connectivity within the DMN and TPN whilst contributing towards their overall anti-correlation.

There were wide-ranging significant differences in functional connectivity between resting-state and task-performing individuals. This was expected due to the significantly altered functional architecture of the brain during task-performing states compared to resting-states (Elton et al. 2015; Goparaju et al. 2014; Thompson et al. 2013). The significance of these functional connectivity differences is beyond the scope of this study. However, they confirm work done by previous studies highlighting functional connectivity differences between resting-state and task-performing individuals. Noteworthy for this paper is the decrease in the number of functional connectivity differences between resting-state and task-performing scans after regression of the QPP. Many of the functional connectivity differences seen between resting-state and task-performing individuals were diminished significantly once the QPP was regressed. This suggests that the different brain states could partially be a result of QPP activity.

### 4.5 Implications for fMRI

Resting-state fMRI is popular for patient groups as it does not require the performance of a task, reducing the need for active patient cooperation. Alterations in functional connectivity have been observed in numerous neurological and psychiatric disorders, especially in the DMN. The differences in functional connectivity tend to be interpreted in terms of network interactions (i.e., a brain region is hypo-connected, or modularity is decreased (Mohan et al. 2016)). However, the presence of QPPs suggests a complementary interpretation where activity within and between networks is coordinated by a non-localized mechanism that simultaneously modulates activity in large swaths of the brain. Thus, the disruption of functional connectivity could at least in part reflect dysfunction of the process that produces QPPs. A recent paper shows that different brainstem nuclei are linked to activity in the DMN and TPN (Bär et al. 2016). The QPP could arise from coordinated input from these neuromodulatory regions, a hypothesis supported by preliminary findings that QPPs are weaker in rats with diminished locus coeruleus activity (Abbas et al. 2018b). Several neurological disorders such as Alzheimer’s Disease and Parkinson’s Disease exhibit early degeneration of neuromodulatory nuclei, which could then account for the disrupted functional connectivity that is observed in those individuals.

Besides implications for clinical functional connectivity studies, the strong contribution of QPPs to functional connectivity also affects the interpretation of more basic neuroscience studies. If QPPs are related to neuromodulatory input and arousal, changes in functional connectivity observed during task performance may be tied to increased arousal during difficult tasks and lower levels of arousal during less difficult tasks. This complicates the use of functional connectivity to understand how the large-scale networks of the brain are reorganized for optimal task performance.

### 4.6 Limitations

While the pattern-finding algorithm depends on a few parameters that must be chosen by a user, a substantial body of work has shown that QPPs can be reliably detected in multiple species, under different physiological conditions, and by using several variations of the basic pattern-finding algorithm. Hence, QPP detection appears quite robust. This study builds upon previous work to examine the contribution of the QPPs to functional connectivity using regression.

The use of regression to minimize the contribution of QPPs fundamentally assumes that QPPs are additive to the remaining BOLD signal. Multi-modal experiments in rodents support this assumption: QPPs are more closely linked to infra-slow activity while dynamic measures of BOLD correlation are more reflective of higher frequency activity (Thompson et al. 2014a), and no phase-amplitude coupling was consistently observed between the infra-slow activity and higher frequencies (Thompson et al. 2014b). The lack of phase-amplitude coupling does not rule out other types of interactions such as phase-phase coupling or amplification, but it suggests that treating QPPs as an additive signal is a reasonable first approximation. Further work using animal models is needed where neural recordings can provide a ‘ground truth’ comparison.

Furthermore, while the QPP can be generally described as involving the DMN and TPN, it is clear from this study that the precise brain areas involved can vary across cognitive conditions. The spatiotemporal differences seen between the QPPs during rest and task performance in this study may be specifically reflective of a working memory task, or they may reflect a general shift between task performance and rest. Participants were requested to keep their eyes fixated and stay awake for both rest and task conditions, so the differences in QPPs should not be related to state differences associated with eye closure. However, it is well-established that participants do tend to fall asleep during resting-state scans. In our study, only the first 5 minutes of 15-minute resting-state scans were used, which hopefully reduces the effect sleepiness or drowsiness could have had on our results and conclusions. Further work will be necessary to determine if different tasks involving different brain regions may affect the QPP in a unique way.

Finally, it is important to mention our decision to implement global signal regression during preprocessing. Yousefi et al. (2018) demonstrated that global signal regression reduces variability in QPPs acquired from different subjects. In the study, subjects were divided into two groups; those with low levels of global signal fluctuation and those with high levels of global signal fluctuation. Subjects with low levels of global signal fluctuation showed a QPP demonstrating anti-correlated network activity, as has been described in this paper. Subjects with high levels of global signal fluctuation showed that the global signal had an additive effect on the QPP: Though the observed spatial distribution of the pattern and its frequency of occurrence was relatively unchanged, the whole-brain global changes in BOLD signal obscured the underlying pattern. When global signal regression was conducted on individuals with high levels of global signal fluctuation, their QPPs aligned with the QPPs of individuals with low levels of global signal fluctuation.

For this paper, the aim was to understand the effects of QPP regression on functional connectivity in the brain. If global signal had not been regressed from the functional scans, it could have served as a confounding factor in the subsequent analysis. Depending on the levels of global signal fluctuation in each subject, the spatiotemporal pattern observed in QPPs would have varied and their regression would have affected static functional connectivity differently across subjects. Hence, for a study investigating the effect of QPP regression on functional connectivity, we believe global signal regression in all functional scans was the appropriate decision, especially given that there are existing studies that demonstrate the effects of global signal regression on functional connectivity (Murphy & Fox 2017).

## 5 Conclusion

Quasi-periodic patterns can be detected in both resting-state and task-performing individuals, with the task influencing the spatiotemporal pattern seen within the QPP as well as the strength and frequency of its occurrence. Removal of QPPs from functional scans through linear regression leads to significant changes in functional connectivity, especially within the DMN and TPN. This suggests that QPPs are relevant to healthy brain function and may account for changes in connectivity in certain patient groups. The findings also suggest that infra-slow electrical activity reflected by QPPs may play a role in the organization of network activity within the brain.

## Acknowledgements

A sincere thank you Michael Borich, Helen Mayberg, Paul Garcia, David Weinshenker, Behnaz Yousefi, Wenju Pan, and Xiaodi Zhang for participating in lively discussions regarding this work. This work was supported by National Science Foundation BCS INSPIRE 1533260, National Institutes of Health R01NS078095, National Institutes of Health 1R01MH111416-01, and the ISMRM Research Exchange Program (granted to MB). Data were provided by the Human Connectome Project, WU-Minn Consortium (Principal Investigators: David Van Essen and Kamil Ugurbil; 1U54MH091657) funded by the 16 NIH Institutes and Centers that support the NIH Blueprint for Neuroscience Research; and by the McDonnell Center for Systems Neuroscience at Washington University.

---

**Supplementary Figure 1:**
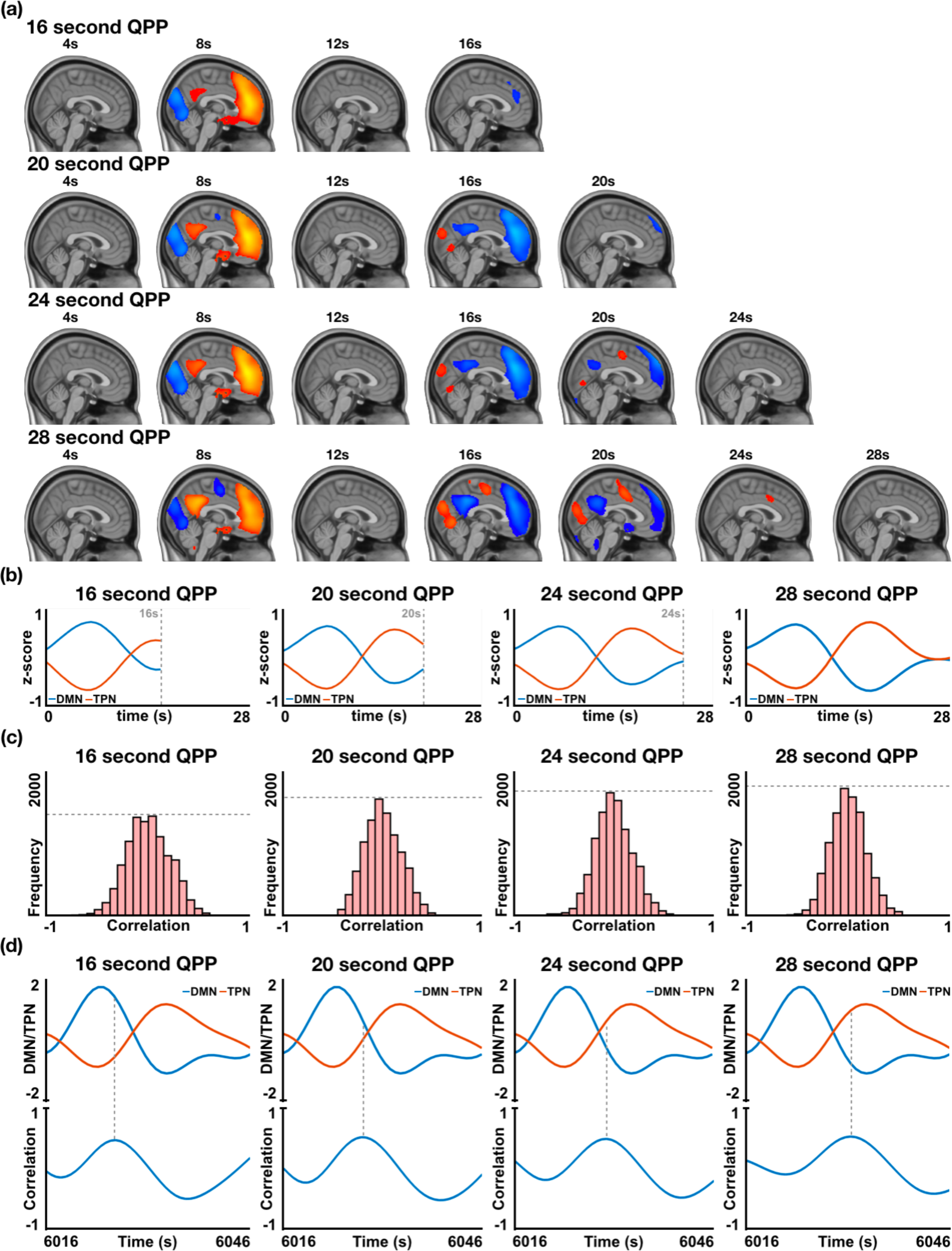
In resting-state individuals, the effect of inputting different window lengths when searching for QPPs using the spatiotemporal pattern-finding algorithm, with the purpose of determining an apt QPP length to use for this study. **(a)** Spatiotemporal pattern of four QPPs with lengths 16s, 20s, 24s, and 28s (top to bottom). **(b)** Timecourses of the DMN and TPN during each of the QPPs. As can be observed, 16s is too short of a length to capture a complete transition between DMN and TPN dominance, hence this length can be discarded in our search for an apt QPP length. **(c)** Histogram of the cumulative sliding correlation of the QPP with the 25 concatenated scans it was acquired from. Of the three QPP lengths remaining, the strongest correlation (as determined by the widest distribution) is shown by the 20s QPP. With increasing QPP length, the strength of the sliding correlation decreases. **(d)** *Top:* Timecourse of the DMN and TPN at an instance in the functional data where a DMN to TPN transition is occurring. *Bottom:* Sliding correlation of the QPP with the functional scan at the same instance where a DMN to TPN transition is occurring. For an ideal QPP length, peaks in the sliding correlation vector should occur in the center of the QPP, which is the exact instance at which the DMN/TPN switch occurs. The 20s and 24s QPPs both pass this test with similar performance. Given the tests carried out in (b), (c), and (d), the 20s QPP stands out as an ideal length of a QPP in resting-state individuals.

**Supplementary Figure 2:**
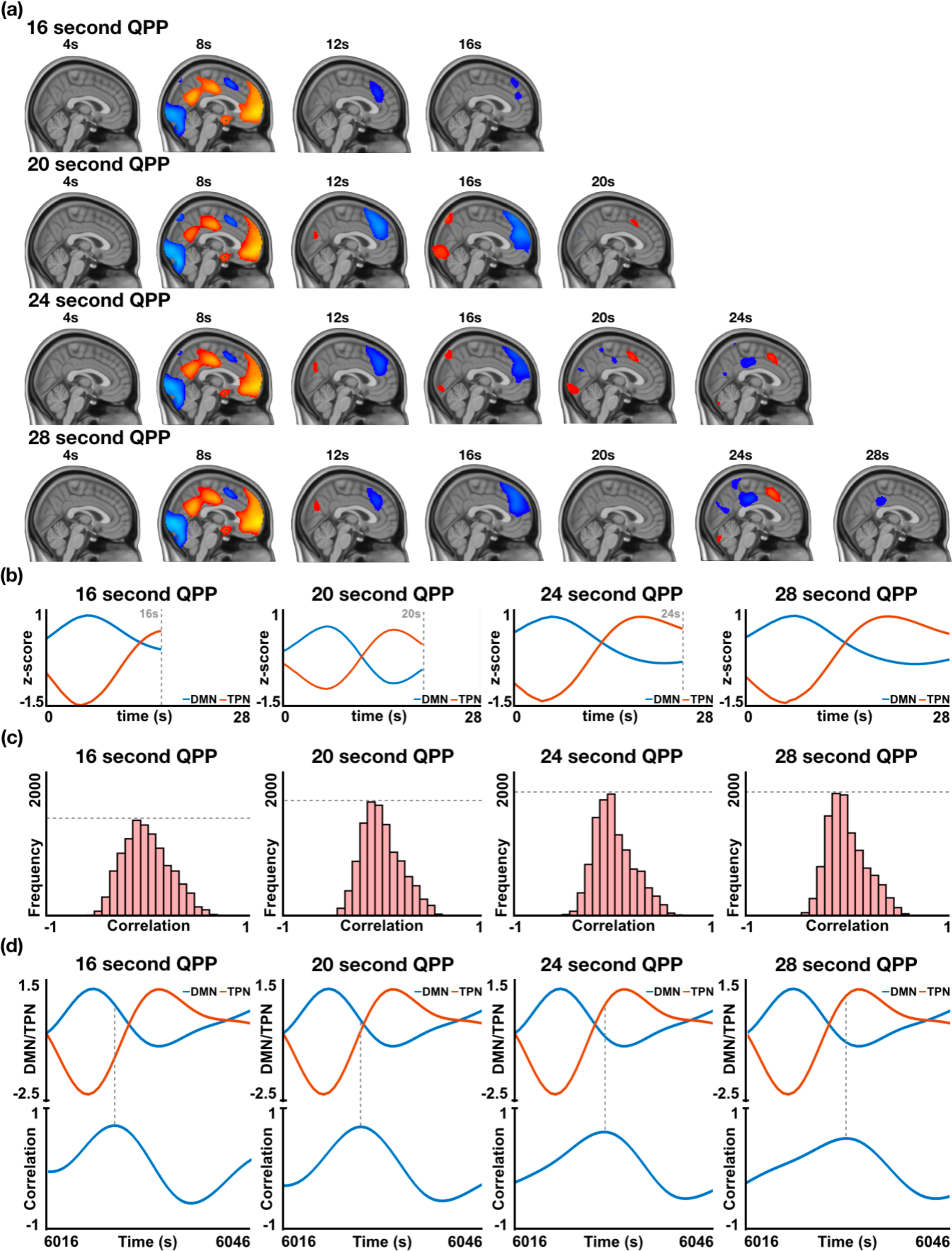
In task-performing individuals, the effect of inputting different window lengths when searching for QPPs using the spatiotemporal pattern-finding algorithm, with the purpose of determining an apt QPP length to use for this study. **(a)** Spatiotemporal pattern of four QPPs with lengths 16s, 20s, 24s, and 28s (top to bottom). **(b)** Timecourses of the DMN and TPN during each of the QPPs. As can be observed, 16s is too short of a length to capture a complete transition between DMN and TPN dominance, hence this length can be discarded in our search for an apt QPP length. **(c)** Histogram of the cumulative sliding correlation of the QPP with the 25 concatenated scans it was acquired from. Of the three QPP lengths remaining, the strongest correlation (as determined by the widest distribution) is shown by the 20s QPP. With increasing QPP length, the strength of the sliding correlation decreases. **(d)** *Top:* Timecourse of the DMN and TPN at an instance in the functional data where a DMN to TPN transition is occurring. *Bottom:* Sliding correlation of the QPP with the functional scan at the same instance where a DMN to TPN transition is occurring. For an ideal QPP length, peaks in the sliding correlation vector should occur in the center of the QPP, which is the exact instance at which the DMN/TPN switch occurs. The 20s QPP passes this test with the best performance. Given the tests carried out in (b), (c), and (d), the 20s QPP stands out as an ideal length of a QPP in task-performing individuals.

**Supplementary Figure 3.**
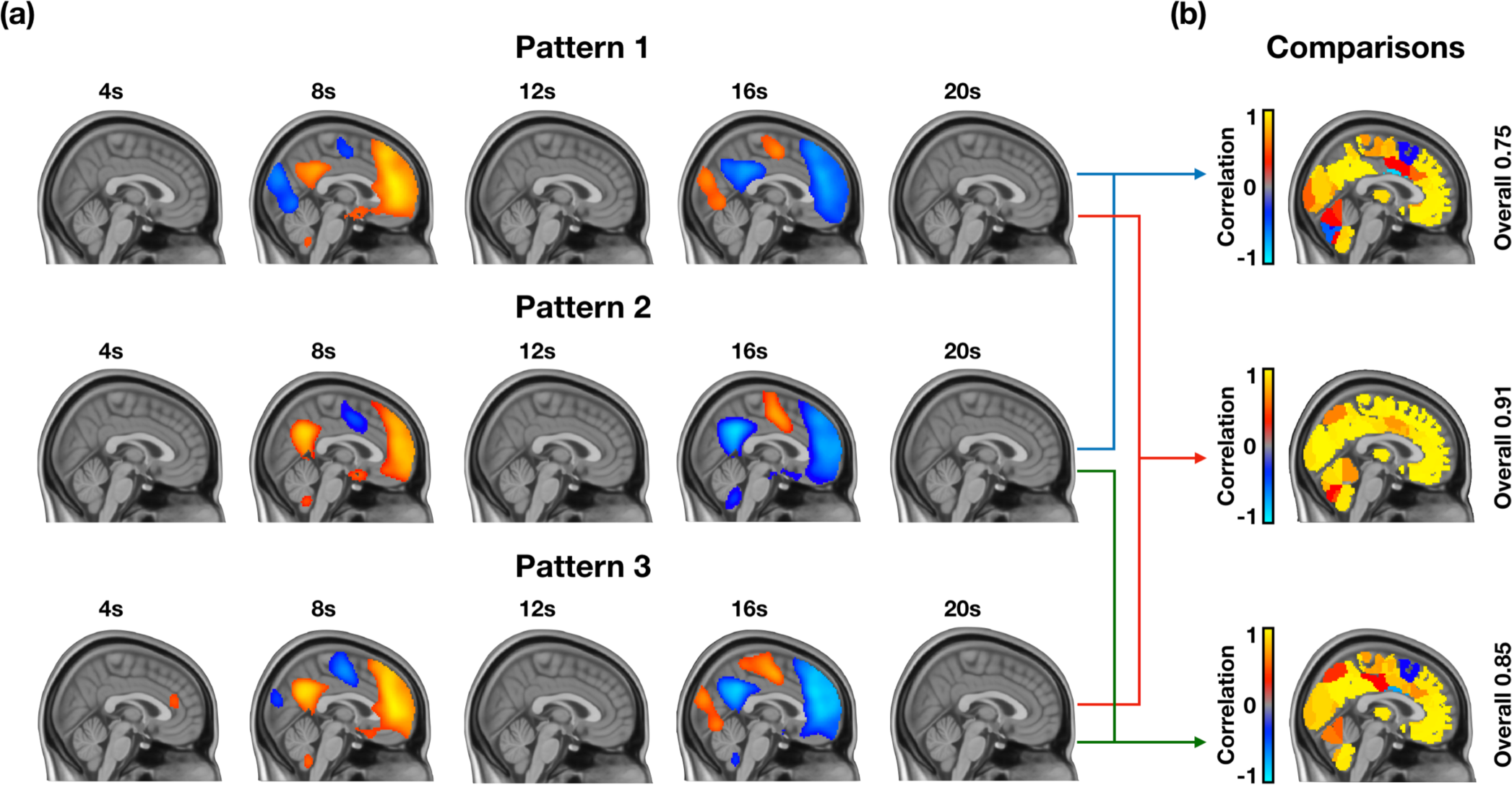
Lack of variability in QPPs acquired from three different groups of 25 individuals. **(a)** Quasi-periodic patterns from three different groups of 25 individuals all show a DMN-to-TPN transition in BOLD activity. **(b)** Correlation between the timecourses of 273 ROIs within each QPP shows that most regions are highly correlated across QPPs. This shows that the QPPs used for the groups were representative of the group and that concatenation of data from 25 individuals was sufficient in acquiring a QPP for that group. The analysis for this figure was done in resting-state individuals. A similar analysis for task-performing individuals is done in Supplementary Figure 4.

**Supplementary Figure 4:**
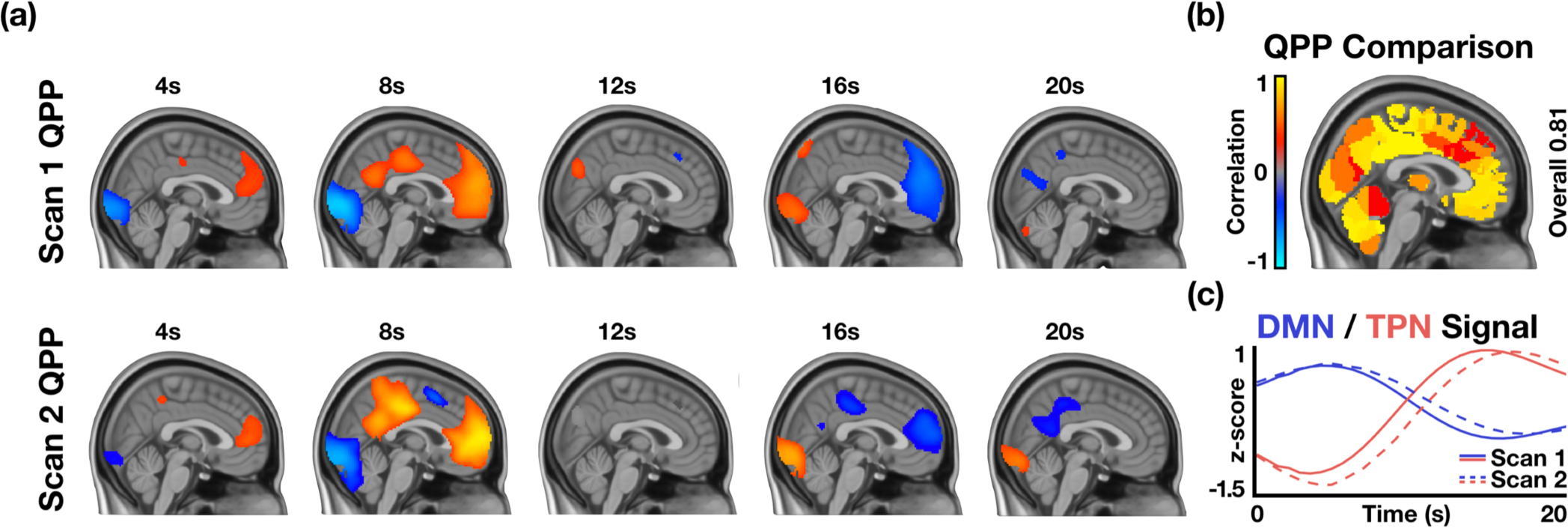
Lack of variability in patterns acquired from the first task-performing functional scan and second task-performing functional scan. **(a)** *Top:* Quasi-periodic pattern acquired by concatenating the first task-performing scan from 25 subjects and applying the spatiotemporal pattern-finding algorithm. *Bottom:* Quasi-periodic pattern acquired by concatenating the second task-performing scan from the same 25 subjects and applying the spatiotemporal pattern-finding algorithm. **(b)** Correlation between the timecourses of 274 ROIs within each QPP shows that most regions are highly correlated across the two QPPs. **(c)** DMN and TPN timecourses in each of the two QPPs show that network behavior is also similar in both QPPs. The analysis in this figure shows that the acquired task-performing scan is representative of the group and that the QPP acquired from the first task-performing scan is similar to the one acquired from the second task-performing scan.

**Supplementary Figure 5:**
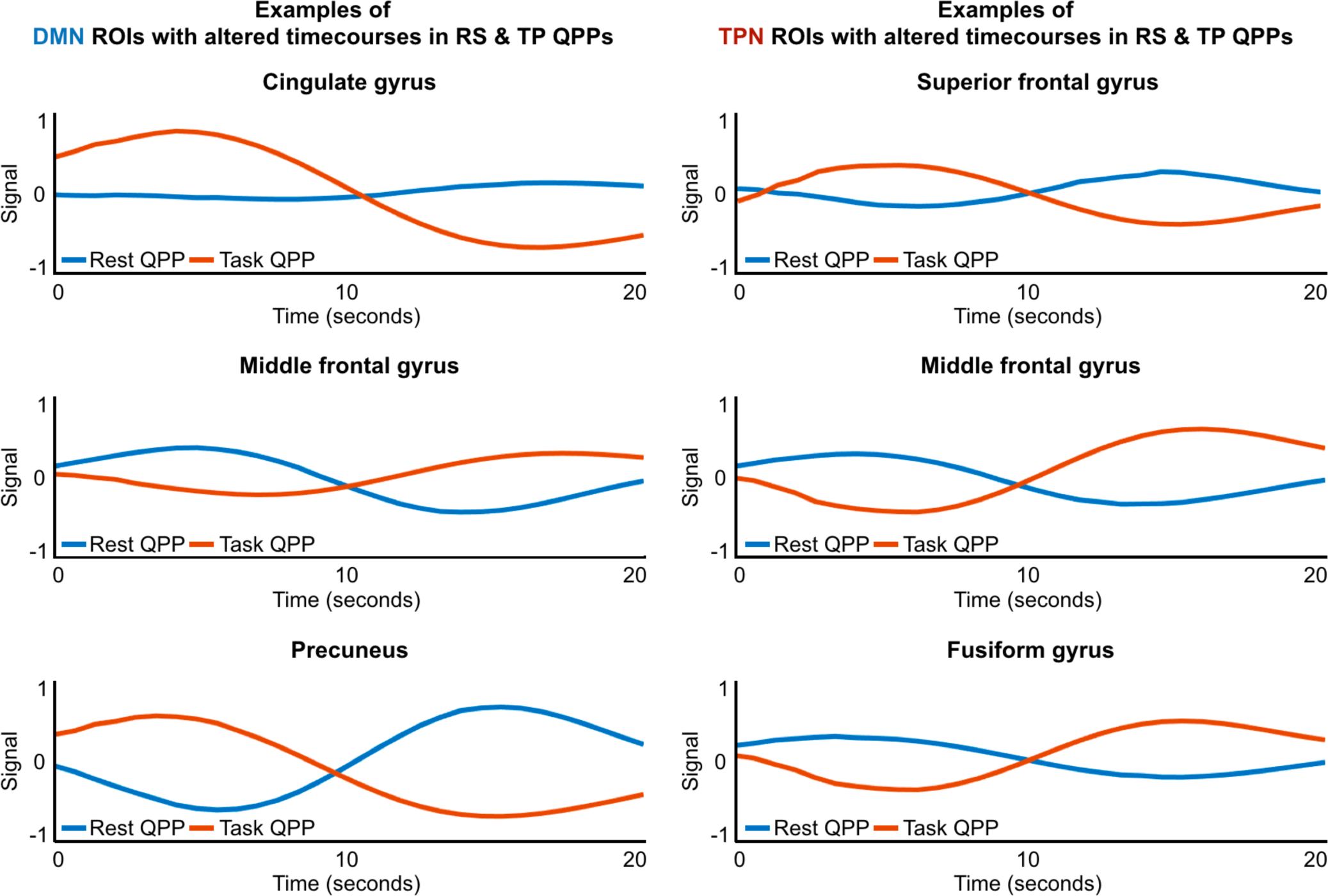
Timecourses of example ROIs that were identified as behaving differently in the resting-state QPP versus the task-performing QPP. Blue lines show how the ROI behaved during the resting-state QPP while red lines show how the ROI behaved during the task-performing QPP.

**Supplementary Figure 6:**
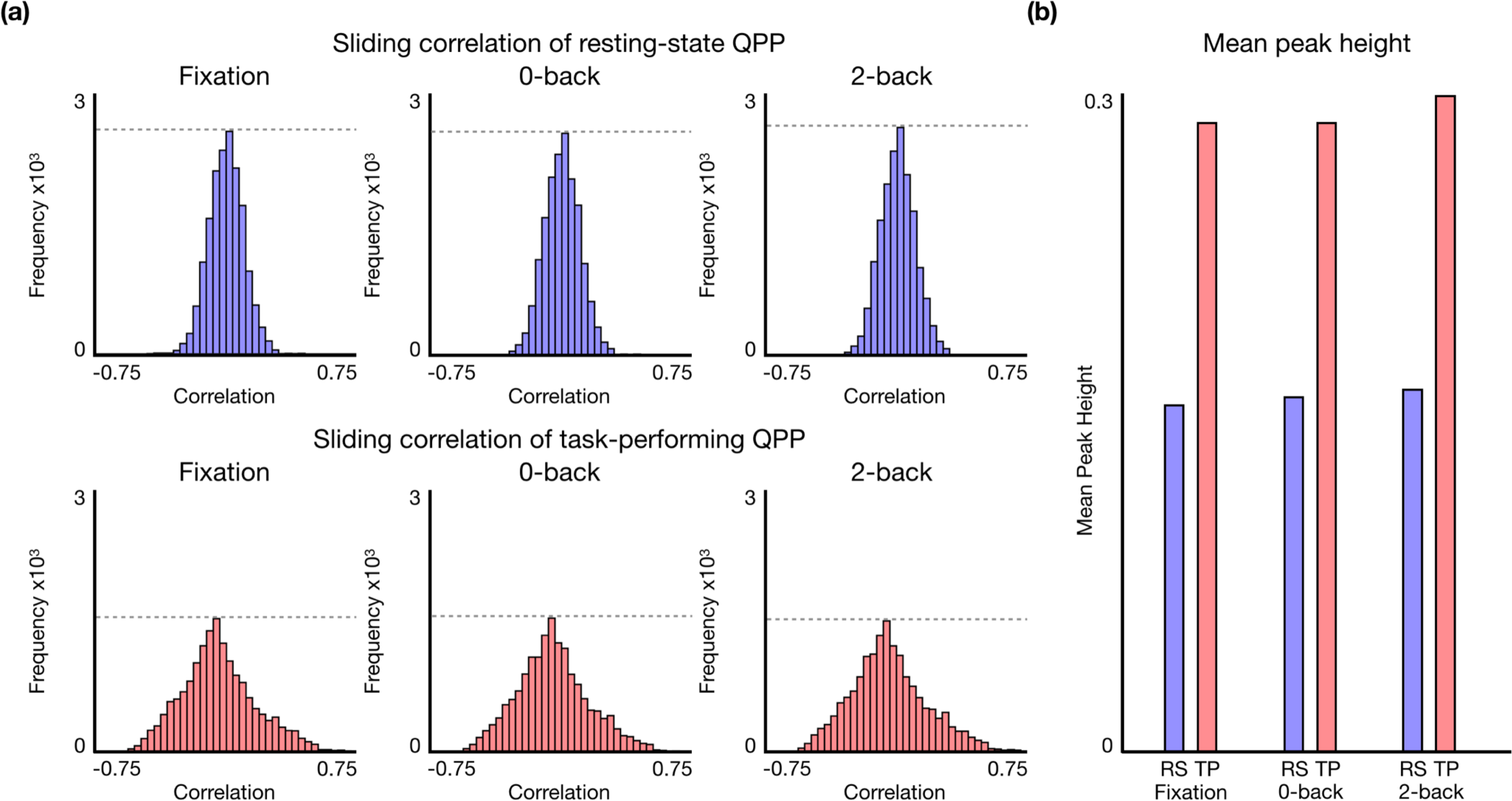
Effects of fixation, 0-back, and 2-back blocks on the occurrence of resting-state and task-performing QPPs in task-performing scans. The sliding correlation vectors of each of the QPPs were separated into areas that corresponded with the 15s fixation blocks and 25s 0-back and 2-back blocks in each scan. **(a)** *Top:* Histograms of the cumulative sliding correlation of the resting-state QPP with instances in all the task-performing scans that corresponded with fixation (left), the 0-back working memory task (middle), and the 2-back working memory task (right). There was no significant difference in the distribution of the sliding correlation vector of the resting-state QPP in each of the blocks. *Bottom:* Histograms of the cumulative sliding correlation of the task-performing QPP with instances in all the task-performing scans that corresponded with fixation (left), the 0-back working memory task (middle), and the 2-back working memory task (right). There was no significant difference in the distribution of the sliding correlation vector of the task-performing QPP in each of the blocks. As previously shown in Figure 2, the presence of the task-performing QPP was stronger in the task-performing scans compared to the resting-state QPP. **(b)** Mean correlation strength for all peaks > 0.1 in the sliding correlation vectors of the resting-state and task-performing QPPs in each of the blocks during all the task-performing scans. There was no significant difference in the mean peak height of either the resting-state or task-performing QPPs across the blocks in the task-performing scans. As previously shown in Figure 2, the mean peak heights of the task-performing QPP in task-performing scans is greater than those of the resting-state QPP.

**Supplementary Figure 7.**
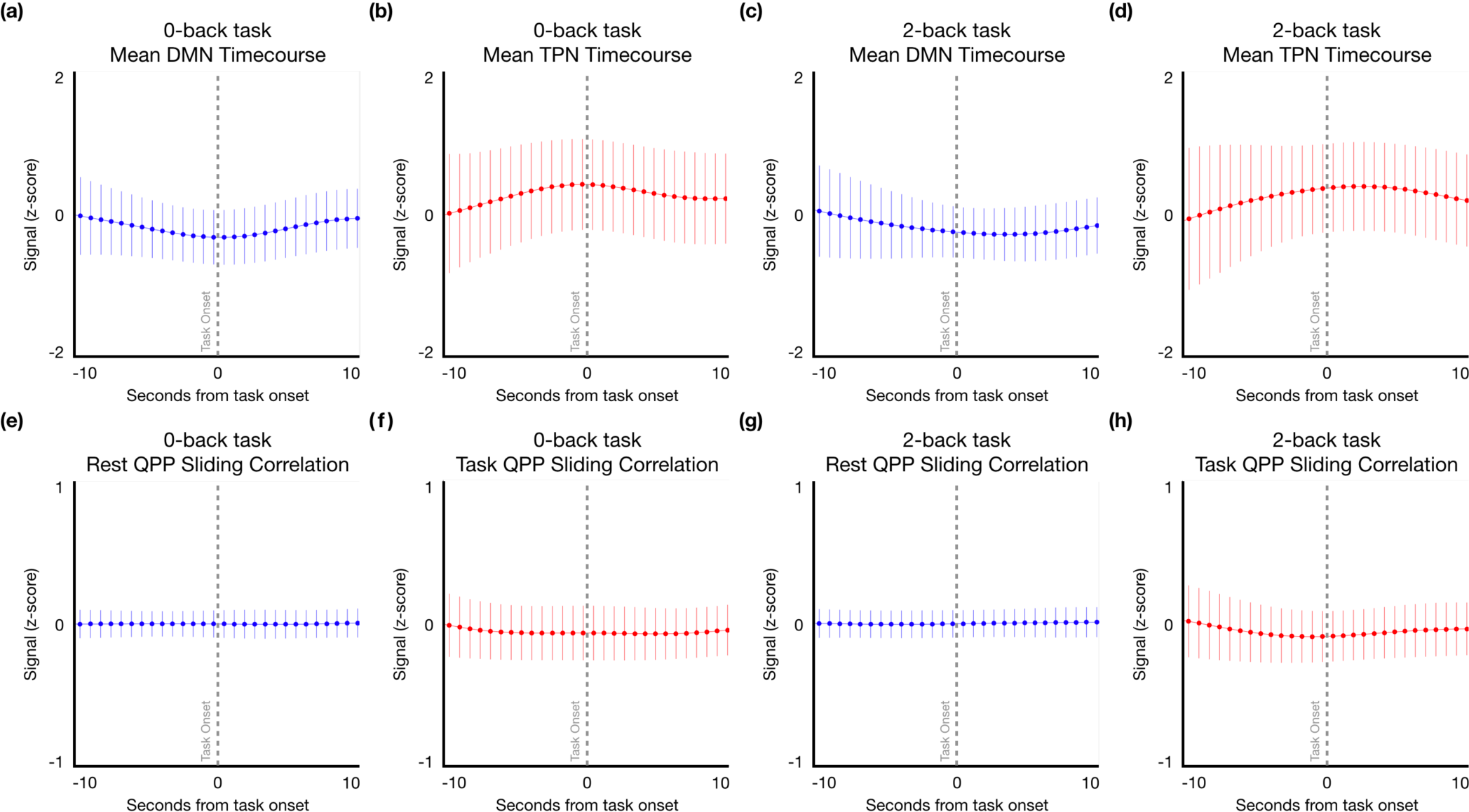
Effect on task stimulus onset on the DMN timecourse, TPN timecourse, and sliding window correlation of the resting-state and task-performing QPPs in task-performing scans. With four 0-back task blocks and 2-back task blocks each in every functional scan, there were a total of 800 task onsets to record. **(a)** Mean DMN timecourse during all 0-back task onsets. **(b)** Mean TPN timecourse during all 0-back task onsets. **(c)** Mean DMN timecourse during all 2-back task onsets. **(d)** Mean TPN timecourse during all 2-back task onsets. **(e)** Mean sliding correlation of the resting-state QPP during all 0-back task onsets **(f)** Mean sliding correlation of the resting-state QPP during all 2-back task onsets. **(g)** Mean sliding correlation of the task-performing QPP during all 0-back task onsets. **(h)** Mean sliding correlation of the task-performing QPP during all 2-back task onsets. As can be deduced from the plots, the task onsets did not have a noticeable effect on either the timecourses of the DMN and TPN or the sliding correlation vectors of the resting-state and task-performing QPPs.

## References

Abbas, A. Bassil, Y., Keilholz, S. (2018a). Quasi-periodic patterns contribute to functional connectivity differences in individuals with ADHD. In: Organization for Human Brain Mapping Annual Meeting.

Abbas, A., Langley, J., Howell, L., & Keilholz, S. (2016a). Quasiperiodic patterns vary in frequency between anesthetized and awake monkeys. In: Resting State Brain Connectivity Biennial Conference, p. 141.

Abbas, A., Nezafati, M., Thomas, I., Keilholz, S. (2018b). Quasiperiodic patterns in BOLD fMRI reflect neuromodulatory input. In: International Society for Magnetic Resonance in Medicine 26th Annual Meeting. Abstract #8422.

Andrews-Hanna, J. R., Snyder, A. Z., Vincent, J. L., Lustig, C., Head, D., Raichle, M. E., & Buckner, R. L. (2007). Disruption of Large-Scale Brain Systems in Advanced Aging. Neuron, 56(5), 924–935. http://doi.org/10.1016/j.neuron.2007.10.038

Asemi, A., Ramaseshan, K., Burgess, A., Diwadkar, V. A., & Bressler, S. L. (2015). Dorsal anterior cingulate cortex modulates supplementary motor area in coordinated unimanual motor behavior. Frontiers in Human Neuroscience, 9, 309. http://doi.org/10.3389/fnhum.2015.00309

Bär, K.-J., la Cruz, de, F., Schumann, A., Koehler, S., Sauer, H., Critchley, H., & Wagner, G. (2016). Functional connectivity and network analysis of midbrain and brainstem nuclei. NeuroImage, 134, 53–63. http://doi.org/10.1016/j.neuroimage.2016.03.071

Barch, D. M., Burgess, G. C., Harms, M. P., Petersen, S. E., Schlaggar, B. L., Corbetta, M., et al. (2013). Function in the human connectome: Task-fMRI and individual differences in behavior. NeuroImage, 80(C), 169–189. http://doi.org/10.1016/j.neuroimage.2013.05.033

Belloy, M. E., Naeyaert, M., Abbas, A., Shah, D., Vanreusel, V., van Audekerke, J., et al. (2018). Dynamic resting state fMRI analysis in mice reveals a set of Quasi-Periodic Patterns and illustrates their relationship with the global signal. NeuroImage, 1–22. http://doi.org/10.1016/j.neuroimage.2018.01.075

Benjamini, Y., & Yekutieli, D. (2001). The Control of the False Discovery Rate in Multiple Testing Under Dependency. The Annals of Statistics, 29(4), 1165–1188.

Biswal, B., Zerrin Yetkin, F., Haughton, V. M., & Hyde, J. S. (1995). Functional connectivity in the motor cortex of resting human brain using echo-planar MRI. Magnetic Resonance in Medicine, 34(4), 537–541. http://doi.org/10.1002/mrm.1910340409

Buzsáki, G. (2006). Rhythms of the Brain (pp. 1–465). Oxford University Press, Inc.

Canolty, R. T., & Knight, R. T. (2010). The functional role of cross-frequency coupling. Trends in Cognitive Sciences, 14(11), 506–515. http://doi.org/10.1016/j.tics.2010.09.001

Chang, C., & Glover, G. H. (2010). Time–frequency dynamics of resting-state brain connectivity measured with fMRI. NeuroImage, 50(1), 81–98. http://doi.org/10.1016/j.neuroimage.2009.12.011

Chen, L., Vu, A. T., Xu, J., Moeller, S., Ugurbil, K., Yacoub, E., & Feinberg, D. A. (2015). Evaluation of highly accelerated simultaneous multi-slice EPI for fMRI. NeuroImage, 104(C), 452–459. http://doi.org/10.1016/j.neuroimage.2014.10.027

Choi, J., Jeong, B., Lee, S. W., & Go, H.-J. (2013). Aberrant Development of Functional Connectivity among Resting State-Related Functional Networks in Medication-Naïve ADHD Children. Plos One, 8(12), e83516–11. http://doi.org/10.1371/journal.pone.0083516

Cole, M. W., Ito, T., Bassett, D. S., & Schultz, D. H. (2016). Activity flow over resting-state networks shapes cognitive task activations. Nature Neuroscience, 19(12), 1718–1726. http://doi.org/10.1038/nn.4406

Elton, A., & Gao, W. (2015). Task-related modulation of functional connectivity variability and its behavioral correlations. Human Brain Mapping, 36(8), 3260–3272. http://doi.org/10.1002/hbm.22847

Fan, L., Li, H., Zhuo, J., Zhang, Y., Wang, J., Chen, L., et al. (2016). The Human Brainnetome Atlas: A New Brain Atlas Based on Connectional Architecture. Cerebral Cortex, 26(8), 3508–3526. http://doi.org/10.1093/cercor/bhw157

Feinberg, D. A., Moeller, S., Smith, S. M., Auerbach, E., Ramanna, S., Glasser, M. F., et al. (2010). Multiplexed Echo Planar Imaging for Sub-Second Whole Brain FMRI and Fast Diffusion Imaging. Plos One, 5(12), e15710–11. http://doi.org/10.1371/journal.pone.0015710

Foster, B. L., He, B. J., Honey, C. J., Jerbi, K., Maier, A., & Saalmann, Y. B. (2016). Spontaneous Neural Dynamics and Multi-scale Network Organization. Frontiers in Systems Neuroscience, 10(309), 7. http://doi.org/10.3389/fnsys.2016.00007

Fox, M. D., Snyder, A. Z., Vincent, J. L., Corbetta, M., Van Essen, D. C., & Raichle, M. E. (2005). The human brain is intrinsically organized into dynamic, anticorrelated functional networks. PNAS, 102(27), 9673–9678. http://doi.org/10.1073/pnas.0504136102

Gillebert, C. R., & Mantini, D. (2013). Functional connectivity in the normal and injured brain. The Neuroscientist : a Review. The Neuroscientist, 19(5), 509–522. http://doi.org/10.1177/1073858412463168

Goparaju, B., Rana, K. D., Calabro, F. J., & Vaina, L. M. (2014). A computational study of whole-brain connectivity in resting state and task fMRI. Medical Science Monitor, 20, 1024–1042. http://doi.org/10.12659/MSM.891142

Grimm, S., Boesiger, P., Beck, J., Schuepbach, D., Bermpohl, F., Walter, M., et al. (2008). Altered Negative BOLD Responses in the Default-Mode Network during Emotion Processing in Depressed Subjects. Neuropsychopharmacology, 34(4), 932–943. http://doi.org/10.1038/npp.2008.81

Grooms, J. K., Thompson, G. J., Pan, W.-J., Billings, J., Schumacher, E. H., Epstein, C. M., & Keilholz, S. D. (2017). Infraslow Electroencephalographic and Dynamic Resting State Network Activity. Brain Connectivity, 7(5), 265–280. http://doi.org/10.1089/brain.2017.0492

Hampson, M., Driesen, N., Roth, J. K., Gore, J. C., & Constable, R. T. (2010). Functional connectivity between task-positive and task-negative brain areas and its relation to working memory performance. Magnetic Resonance Imaging, 28(8), 1051–1057. http://doi.org/10.1016/j.mri.2010.03.021

He, B. J., Snyder, A. Z., Zempel, J. M., Smyth, M. D., & Raichle, M. E. (2008). Electrophysiological correlates of the brain’s intrinsic large-scale functional architecture. PNAS, 105(41), 16039–16044.

Hiltunen, T., Kantola, J., Abou Elseoud, A., Lepola, P., Suominen, K., Starck, T., et al. (2014). Infra-Slow EEG Fluctuations Are Correlated with Resting-State Network Dynamics in fMRI. Journal of Neuroscience, 34(2), 356–362. http://doi.org/10.1523/JNEUROSCI.0276-13.2014

Hoekzema, E., Carmona, S., Ramos-Quiroga, J. A., Richarte Fernández, V., Bosch, R., Soliva, J. C., et al. (2013). An independent components and functional connectivity analysis of resting state fMRI data points to neural network dysregulation in adult ADHD. Human Brain Mapping, 35(4), 1261–1272. http://doi.org/10.1002/hbm.22250

Holt, D. J., Cassidy, B. S., Andrews-Hanna, J. R., Lee, S. M., Coombs, G., Goff, D. C., et al. (2011). An Anterior-to-Posterior Shift in Midline Cortical Activity in Schizophrenia During Self-Reflection. Biol Psychiatry, 69(5), 415–423. http://doi.org/10.1016/j.biopsych.2010.10.003

Hutchison, R. M., Womelsdorf, T., Allen, E. A., Bandettini, P. A., Calhoun, V. D., Corbetta, M., et al. (2013). Dynamic functional connectivity: Promise, issues, and interpretations. NeuroImage, 80, 360–378. http://doi.org/10.1016/j.neuroimage.2013.05.079

Jenkinson, M., & Smith, S. (2001). A global optimisation method for robust affine registration of brain images. Medical Image Analysis, 5(2), 143–156.

Jenkinson, M., Bannister, P., Brady, M., & Smith, S. (2002). Improved optimization for the robust and accurate linear registration and motion correction of brain images. NeuroImage, 17(2), 825–841.

Jenkinson, M., Beckmann, C. F., Behrens, T. E. J., Woolrich, M. W., & Smith, S. M. (2012). FSL. NeuroImage, 62(2), 782–790. http://doi.org/10.1016/j.neuroimage.2011.09.015

Kelly, A. M. C., Uddin, L. Q., Biswal, B. B., Castellanos, F. X., & Milham, M. P. (2008). Competition between functional brain networks mediates behavioral variability. NeuroImage, 39(1), 527–537. http://doi.org/10.1016/j.neuroimage.2007.08.008

Liddle, E. B., Hollis, C., Batty, M. J., Groom, M. J., Totman, J. J., Liotti, M., et al. (2010). Task-related default mode network modulation and inhibitory control in ADHD: effects of motivation and methylphenidate. Journal of Child Psychology and Psychiatry, 52(7), 761–771. http://doi.org/10.1111/j.1469-7610.2010.02333.x

Lin, H.-Y., & Gau, S. S.-F. (2016). Atomoxetine Treatment Strengthens an Anti-Correlated Relationship between Functional Brain Networks in Medication-Naïve Adults with Attention-Deficit Hyperactivity Disorder: A Randomized Double-Blind Placebo-Controlled Clinical Trial. International Journal of Neuropsychopharmacology, 19(3), pyv094–15. http://doi.org/10.1093/ijnp/pyv094

Majeed, W., Magnuson, M., & Keilholz, S. D. (2009). Spatiotemporal dynamics of low frequency fluctuations in BOLD fMRI of the rat. Journal of Magnetic Resonance Imaging, 30(2), 384–393. http://doi.org/10.1002/jmri.21848

Majeed, W., Magnuson, M., Hasenkamp, W., Schwarb, H., Schumacher, E. H., Barsalou, L., & Keilholz, S. D. (2011). Spatiotemporal dynamics of low frequency BOLD fluctuations in rats and humans. NeuroImage, 54(2), 1140–1150. http://doi.org/10.1016/j.neuroimage.2010.08.030

Matsui, T., Murakami, T., & Ohki, K. (2016). Transient neuronal coactivations embedded in globally propagating waves underlie resting-state functional connectivity. PNAS, 113(23), 6556–6561. http://doi.org/10.1073/pnas.1521299113

Milchenko, M., & Marcus, D. (2012). Obscuring Surface Anatomy in Volumetric Imaging Data. Neuroinformatics, 11(1), 65–75. http://doi.org/10.1007/s12021-012-9160-3

Mitra, A., Snyder, A. Z., Hacker, C. D., & Raichle, M. E. (2014). Lag structure in resting-state fMRI. Journal of Neurophysiology, 111(11), 2374–2391. http://doi.org/10.1152/jn.00804.2013

Mohan, A., Roberto, A., Mohan, A., Lorenzo, A., Jones, K., Carney, M., et al. (2016). The Significance of the Default Mode Network (DMN) in Neurological and Neuropsychiatric Disorders: A Review. Yale Journal of Biology and Medicine, 89(1), 49–57.

Monto, S., Palva, S., Voipio, J., & Palva, J. M. (2008). Very slow EEG fluctuations predict the dynamics of stimulus detection and oscillation amplitudes in humans. The Journal of Neuroscience, 28(33), 8268–8272. http://doi.org/10.1523/JNEUROSCI.1910-08.2008

Murphy, K., & Fox, M. D. (2017). Towards a consensus regarding global signal regression for resting state functional connectivity MRI. NeuroImage, 154, 169–173. http://doi.org/10.1016/j.neuroimage.2016.11.052

Nir, Y., Mukamel, R., Dinstein, I., Privman, E., Harel, M., Fisch, L., et al. (2008). Interhemispheric correlations of slow spontaneous neuronal fluctuations revealed in human sensory cortex. Nature Neuroscience, 11(9), 1100–1108. http://doi.org/10.1038/nn.2177

Palva, J. M., & Palva, S. (2012). Infra-slow fluctuations in electrophysiological recordings, blood-oxygenation-level-dependent signals, and psychophysical time series. NeuroImage, 62(4), 2201–2211. http://doi.org/10.1016/j.neuroimage.2012.02.060

Pan, W.-J., Thompson, G. J., Magnuson, M. E., Jaeger, D., & Keilholz, S. (2013). Infraslow LFP correlates to resting-state fMRI BOLD signals. NeuroImage, 74(C), 288–297. http://doi.org/10.1016/j.neuroimage.2013.02.035

Park, H. J., & Friston, K. (2013). Structural and Functional Brain Networks: From Connections to Cognition. Science, 342(6158), 1238411–1238411. http://doi.org/10.1126/science.1238411

Pievani, M., Filippini, N., van den Heuvel, M. P., Cappa, S. F., & Frisoni, G. B. (2014). Brain connectivity in neurodegenerative diseases—from phenotype to proteinopathy. Nature Reviews Neurology, 10(11), 620–633. http://doi.org/10.1038/nrneurol.2014.178

Posner, J., Park, C., & Wang, Z. (2014). Connecting the Dots: A Review of Resting Connectivity MRI Studies in Attention-Deficit/Hyperactivity Disorder. Neuropsychology Review, 24(1), 3–15. http://doi.org/10.1007/s11065-014-9251-z

Power, J. D., Cohen, A. L., Nelson, S. M., Wig, G. S., Barnes, K. A., Church, J. A., et al. (2011). Functional Network Organization of the Human Brain. Neuron, 72(4), 665–678. http://doi.org/10.1016/j.neuron.2011.09.006

Raichle, M. E. (2011). The restless brain. Brain Connectivity, 1(1), 3–12. http://doi.org/10.1089/brain.2011.0019

Raichle, M. E. (2015). The Brain’s Default Mode Network. Annual Review of Neuroscience, 38(1), 433–447. http://doi.org/10.1146/annurev-neuro-071013-014030

Setsompop, K., Gagoski, B. A., Polimeni, J. R., Witzel, T., Wedeen, V. J., & Wald, L. L. (2011). Blipped-controlled aliasing in parallel imaging for simultaneous multislice echo planar imaging with reduced g-factor penalty. Magnetic Resonance in Medicine, 67(5), 1210–1224. http://doi.org/10.1002/mrm.23097

Sheline, Y. I., Price, J. L., Yan, Z., & Mintun, M. A. (2010b). Resting-state functional MRI in depression unmasks increased connectivity between networks via the dorsal nexus. PNAS, 107(24), 11020–11025. http://doi.org/10.1073/pnas.1000446107

Sheline, Y. I., Raichle, M. E., Snyder, A. Z., Morris, J. C., Head, D., Wang, S., & Mintun, M. A. (2010a). Amyloid Plaques Disrupt Resting State Default Mode Network Connectivity in Cognitively Normal Elderly. Biol Psychiatry, 67(6), 584–587. http://doi.org/10.1016/j.biopsych.2009.08.024

Sheline, Y., Barch, D. M., Price, J., Rundle, M., Vaishnavi, S. N., Snyder, A. Z., et al. (2009). The default mode network and self-referential processes in depression. PNAS, 106(6), 1942–1947.

Smith, S. M., Vidaurre, D., Beckmann, C. F., Glasser, M. F., Jenkinson, M., Miller, K. L., et al. (2013). Functional connectomics from resting-state fMRI. Trends in Cognitive Sciences, 17(12), 666–682. http://doi.org/10.1016/j.tics.2013.09.016

Thompson, G. J., Magnuson, M. E., Merritt, M. D., Schwarb, H., Pan, W.-J., McKinley, A., et al. (2013). Short-time windows of correlation between large-scale functional brain networks predict vigilance intraindividually and interindividually. Human Brain Mapping, 34(12), 3280–3298. http://doi.org/10.1002/hbm.22140

Thompson, G. J., Pan, W.J., Billings, J. C. W., Grooms, J. K., Shakil, S., Jaeger, D., & Keilholz, S. D. (2014a). Phase-amplitude coupling and infraslow (<1 Hz) frequencies in the rat brain: relationship to resting state fMRI. Frontiers in Integrative Neuroscience, 8, 41. http://doi.org/10.3389/fnint.2014.00041

Thompson, G. J., Pan, W.J., Magnuson, M. E., Jaeger, D., & Keilholz, S. D. (2014b). Quasi-periodic patterns (QPP): large-scale dynamics in resting state fMRI that correlate with local infraslow electrical activity. NeuroImage, 84, 1018–1031. http://doi.org/10.1016/j.neuroimage.2013.09.029

van den Heuvel, M. P., & Hulshoff Pol, H. E. (2010). Specific somatotopic organization of functional connections of the primary motor network during resting state. Human Brain Mapping, 31(4), 631–644. http://doi.org/10.1002/hbm.20893

Van Essen, D. C., Ugurbil, K., Auerbach, E., Barch, D., Behrens, T. E. J., Bucholz, R., et al. (2012). The Human Connectome Project: A data acquisition perspective. NeuroImage, 62(4), 2222–2231. http://doi.org/10.1016/j.neuroimage.2012.02.018

Vincent, J. L., Kahn, I., Snyder, A. Z., Raichle, M. E., & Buckner, R. L. (2008). Evidence for a frontoparietal control system revealed by intrinsic functional connectivity. Journal of Neurophysiology, 100(6), 3328–3342. http://doi.org/10.1152/jn.90355.2008

Yousefi, B., Shin, J., Schumacher, E. H., & Keilholz, S. D. (2018). Quasi-periodic patterns of intrinsic brain activity in individuals and their relationship to global signal. NeuroImage, 167, 297–308. http://doi.org/10.1016/j.neuroimage.2017.11.043

Zhang, D., Snyder, A. Z., Fox, M. D., Sansbury, M. W., Shimony, J. S., & Raichle, M. E. (2008). Intrinsic functional relations between human cerebral cortex and thalamus. Journal of Neurophysiology, 100(4), 1740– 1748. http://doi.org/10.1152/jn.90463.2008

Zhang, H., Wang, S., Xing, J., Liu, B., Ma, Z., Yang, M., Zhang, Z., Teng G. (2009). Detection of PCC functional connectivity characteristics in resting-state fMRI in mild Alzheimer’s disease. Behavioural Brain Research, 197(1), 103–108. http://doi.org/10.1016/j.bbr.2008.08.012

Zhang, Y., Brady, M., & Smith, S. (2001). Segmentation of brain MR images through a hidden Markov random field model and the expectation-maximization algorithm. IEEE Transactions on Medical Imaging, 20(1), 45–57. http://doi.org/10.1109/42.906424

